# The effect of activity, energy use, and species identity on environmental DNA shedding of freshwater fish

**DOI:** 10.1101/2020.10.28.359653

**Authors:** Bettina Thalinger, Andreas Rieder, Anna Teuffenbach, Yannick Pütz, Thorsten Schwerte, Josef Wanzenböck, Michael Traugott

## Abstract

The quantitative measurement of environmental DNA (eDNA) from field-collected water samples is gaining importance for the monitoring of fish communities and populations. The interpretation of these signal strengths depends, among other factors, on the amount of target eDNA shed into the water. However, shedding rates are presumably associated with species-specific traits such as physiology and behavior. Although such differences between juvenile and adult fish have been previously detected, the general impact of movement and energy use in a resting state on eDNA release into the surrounding water remains hardly addressed.

In an aquarium experiment, we compared eDNA shedding between seven fish species occurring in European freshwaters. The investigated salmonids, cyprinids, and sculpin exhibit distinct adaptions to microhabitats, diets, and either solitary or schooling behavior. The fish were housed in aquaria with constant water flow and their activity was measured by snapshots taken every 30 s. Water samples for eDNA analysis were taken every 3 h and energy use was determined in an intermittent flow respirometer. After controlling for the effect of fish mass, our results demonstrate a positive correlation between target eDNA quantities as measured with digital PCR, fish activity, and energy use, as well as species-specific differences. For cyprinids, the model based on data from individual fish was only partly transferable to groups, which exhibited lower activity and higher energy use.

Our findings highlight the importance of fish physiology and behavior for the comparative interpretation of taxon-specific eDNA quantities. Species traits should therefore be incorporated into eDNA-based monitoring and conservation efforts.

## 1. Introduction

The sensitivity, non-invasiveness, and cost-efficiency of environmental DNA (eDNA) based methods have been proven for diverse habitats and species making them powerful new tools for conservation biology and biodiversity assessments (Barnes and Turner, 2016; Deiner et al., 2017; Huerlimann et al., 2020). Regarding the detection of fish species, eDNA-based monitoring outperforms traditional methods such as electrofishing: for example, for the detection of the endangered European weather loach, *Misgurnus fossilis* (Sigsgaard et al., 2015), the assessment of fish communities in Australian streams (McColl-Gausden et al., 2020), and the distribution of brook trout, *Salvelinus fontinalis* in a US watershed (Evans et al., 2017). The manifold successes of eDNA-based species detection lead to a call for more standardization and better reporting practices (Goldberg et al., 2016; Minamoto et al., 2020; Thalinger et al., 2020a) and to an international effort for implementing the technology into routine species monitoring (Leese et al., 2016; Pilliod et al., 2019). Although reporting the presence/absence of particular species is the starting point of these endeavors, a more quantitative interpretation of field-derived eDNA data is key for the general application of this technology.

Different processes influence the distribution of eDNA in space and time and the detection probabilities of species from environmental samples, namely the origin, degradation, suspension, resuspension, and transport of eDNA (Barnes and Turner, 2016; Harrison et al., 2019). The latter processes are directly linked to local hydrology (e.g. flow and substrate type (Shogren et al., 2017; Pont et al., 2018; Thalinger et al., 2020b)) and environmental conditions (e.g. water temperature, pH, UV-radiation (Strickler et al., 2015; Lacoursière-Roussel et al., 2016; Tsuji et al., 2017)). The amount of eDNA in the water column is directly linked to fish biomass and originally, this was confirmed for common carp (*Cyprinus carpio*) in an aquarium trial and experimental ponds (Takahara et al., 2012). In subsequent experiments, the positive relationship was confirmed for a range of freshwater and marine fish species (Evans et al., 2016; Lacoursière-Roussel et al., 2016; Sassoubre et al., 2016; Horiuchi et al., 2019; Jo et al., 2020). However, these results were primarily obtained for individuals at the same life stage.

eDNA is released into the environment in the form of mucus, feces, scales, and gametes (Merkes et al., 2014; Barnes and Turner, 2016; Sassoubre et al., 2016; Bylemans et al., 2017). Under natural conditions, differences in fish physiology, diet and behavior are likely to affect this process and confound the interpretation of eDNA-based results from a water body (Klymus et al., 2015). For perch and eel, Maruyama et al. (2014) and Takeuchi et al. (2019), respectively, found lower mass-specific eDNA shedding rates for adults in comparison to juveniles, which is likely caused by the scaling in metabolic rates, excretion rates, and surface area with body mass (discussed in Yates et al., 2020). However, these findings could not be confirmed in another experiment with a salmonid species (Mizumoto et al., 2018). In general, the metabolic rate and activity differ between fish species due to distinct physiology and behavior with pelagic species being more active and displaying higher resting metabolic rates than benthic species (Johnston et al., 1988; Killen et al., 2010). A stress response characterized by elevated metabolism and activity is frequently hypothesized as underlying cause for spiking eDNA levels at the beginning of aquarium experiments. Furthermore, metabolism and activity could generally explain mismatching quantitative results in studies comparing eDNA levels between species in the same water body (Takahara et al., 2012; Maruyama et al., 2014; Evans et al., 2016).

Here, we investigate the effect of fish activity (i.e. movement), energy use (i.e. oxygen use × oxycaloric factor), and species identity in an aquarium experiment with seven fish species commonly occurring in European rivers and streams (Fig. 1). We hypothesized that higher activity (i.e. movement) leads to higher eDNA concentrations as there is more shearing between the fish surface and the surrounding water, and higher volumes are pumped through the gills due to the elevated oxygen demand. Independent of activity, fish species with higher energy use in a resting state potentially also emit more eDNA. Additionally, the species-specific composition of the constantly renewed cutaneous mucus layer (Ángeles Esteban, 2012) might lead to differences between individual taxa.

**Figure 1:**
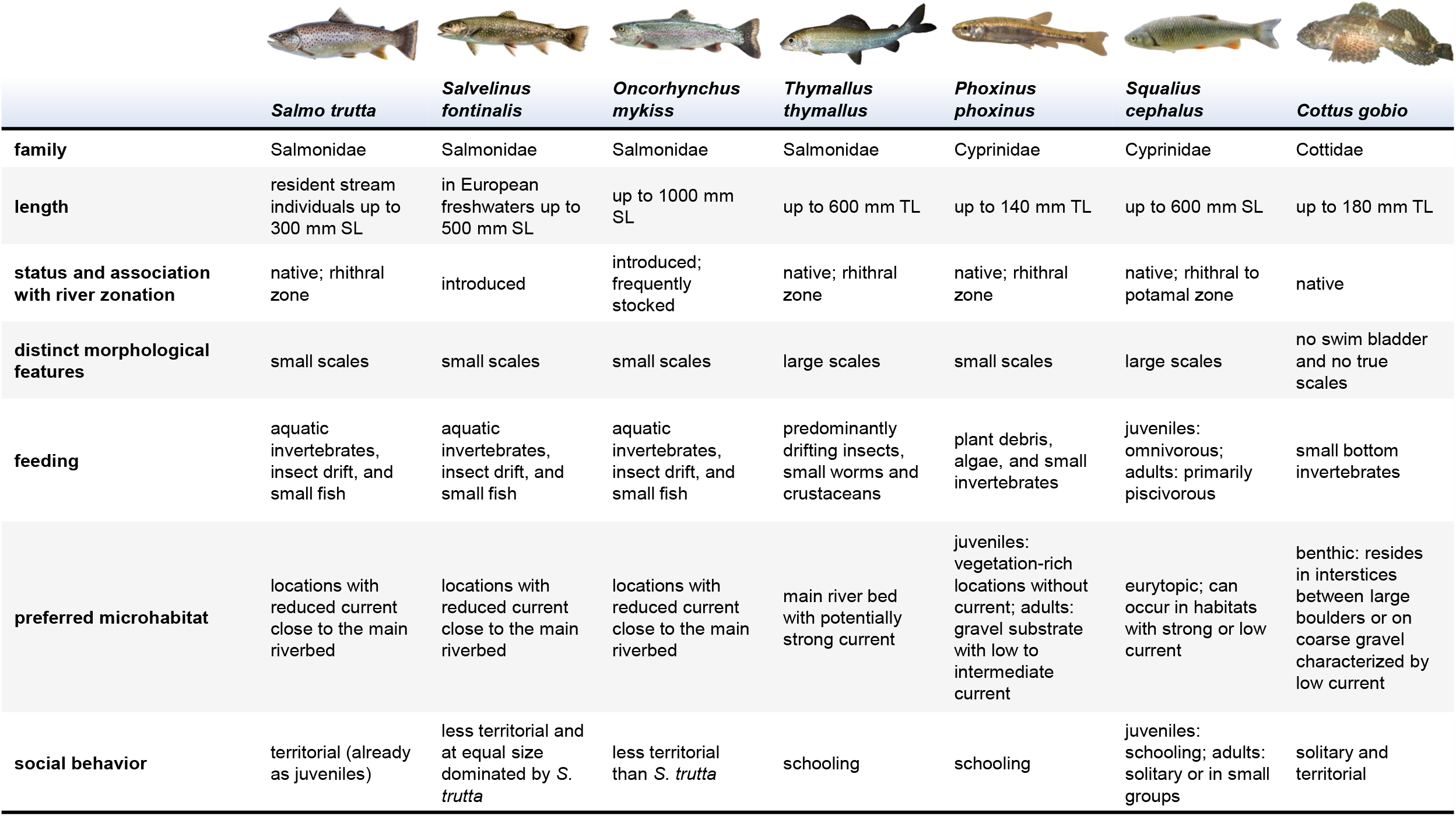
A summary of the morphological and ecological traits of the fish species used in the aquarium experiment. The provided information describes the situation in Central European freshwaters and is not necessarily transferable to other geographic regions. Depending on the source, different maximum fish length measurements were available with “TL” = “total length” measured from the tip of the snout to the longest tip of the caudal fin and “SL” = “standard length” measured from the tip of the snout to the base of the caudal fin. (Muus and Dahlström, 1968; Spindler, 1997; Freyhof and Kottelat, 2007) (https://upload.wikimedia.org/wikipedia/commons/b/bb/CottusGobioSpreadingFins.JPG separated from background; Piet Spaans, CC BY-SA 2.5 https://creativecommons.org/licenses/by-sa/2.5, via Wikimedia Commons; https://upload.wikimedia.org/wikipedia/commons/9/99/Thymallus_thymallus2.jpg separated from background; Gilles San Martin, CC BY-SA 2.0 https://creativecommons.org/licenses/by-sa/2.0, via Wikimedia Commons)

## 2. Materials and methods

### 2.1. Study species

The examined species comprised four salmonids (*Salmo trutta, Salvelinus fontinalis, Oncorhynchus mykiss, Thymallus thymallus*), two cyprinids (*Phoxinus phoxinus, Squalius cephalus*), and one sculpin (*Cottus gobio;* Fig. 1). *Salmo trutta* is a rhithral species, territorial especially in later life stages, and primarily feeds on benthic organisms and insect drift on the surface. *Salvelinus fontinalis* and *O. mykiss* were anthropogenically introduced into European freshwaters and are less territorial than *S. trutta* (Freyhof and Kottelat, 2007). If possible, these three species choose areas with reduced current close to the main riverbed as preferential microhabitat. *Thymallus thymallus* is also a rhithral species, but its scales are larger and adults primarily use the main riverbed (Spindler, 1997; Freyhof and Kottelat, 2007). *Phoxinus phoxinus* is a schooling, small fish species in the rhithral. It feeds on a mixture of plant debris, algae, and small invertebrates. The juveniles prefer vegetation-rich microhabitats without current, while adults switch to gravel substrate with low to intermediate flow. *Squalius cephalus* is eurytopic (rhithral to potamal) and can occur in habitats with strong to low current. Its juveniles are schooling and omnivorous with adults predominantly preying on fish. *Cottus gobio* is a rheophilic and benthic species primarily feeding on small bottom invertebrates. It has no swim bladder and mostly resides in interstices between large boulders or on coarse gravel characterized by low current (Muus and Dahlström, 1968; Spindler, 1997; Freyhof and Kottelat, 2007).

### 2.2. Experimental setup

The aquarium experiment was carried out between 2^nd^ March and 17^th^ July 2017 at the Research Department for Limnology Mondsee of the University of Innsbruck, Austria. The juvenile salmonid individuals were purchased from commercial hatcheries, *P. phoxinus* and *S. cephalus* were caught with permission in Lake Mondsee and *C. gobio* were caught with permission in rivers in Tyrol (Austria). Fish individual sizes were chosen as similar as possible within and between species. As *P. phoxinus* and *C. gobio* are smaller in comparison to the other species (Fig. 1), these individuals were supposedly closer to reproductive maturity. Until the start of the experiment, the fish species were kept separately in aquaria fed with lake water.

In accordance with the regulations of the Austrian Animal Experiment Act (December 28, 2012) (Tierversuchsrechtsänderungsgesetz, part 1, section 1, §1, point 2), and with the Directive 2010/63/EU of the European Parliament and of the Council of the European Union (September 22, 2010) on the protection of animals used for scientific purposes (chapter 1, article 1, point 5a), all fish were reared according to regular agriculture (aquaculture) practice, including the provision of appropriate tank size, sufficient rate of waterflow, natural photoperiod, *ad libitum* food supply, and temperatures within the species’ thermal tolerance range. This ensured that no pain, suffering, distress or lasting harm was inflicted on the animals, confirmed by the fact that mortality rates were low and equal between rearing groups. Based on the legislative provisions above, no ethics approval and no IACUC protocol was required for the experiments performed. In particular the respirometry experiments were discussed with the legislative authorities (Austrian Federal Ministry of Education, Science and Research and the University of Veterinary Medicine, Vienna) and the conclusion was that the assessment of basic metabolism under these conditions (small fish sizes in relatively large chambers) does not incur pain, suffering or distress to the fish and no formal animal experimentation protocol was required.

Five aquaria (60 l) and corresponding plastic lids were used in the experiment, each of which was thoroughly cleaned with sodium hypochlorite (5 %) and then rinsed with tap water (fish-DNA-free) prior to each experimental run (i.e. changing the fish under investigation). The flow-through rate for the tap-water fed aquaria was set to 5.45 l/min to mimic natural conditions and keep eDNA concentrations in the fish tanks constant based on the results of previous test runs (Supplementary Material (SM) 1). The water temperature in the aquaria was stabilized at 15 °C by centrally heating the inflowing water to this temperature. Each tank was further equipped with an air-stone to ensure water mixing. At the start of each experimental run, a water sample (negative control) was taken from one of the aquaria and processed as described below. Then, five fish individuals per species were selected aiming at similar size. Each fish was placed individually in an aquarium using DNA-free fishnets (Fig. 2). For *P. phoxinus* and *S. cephalus*, the experiment was carried out twice: once with individual fish, and once with groups of three fish per aquarium. The day before the experiment and for its duration, the respective fish were not fed to avoid contamination by fish feed and minimize the effects of defecation. Each run started with one day of familiarization.

**Figure 2:**
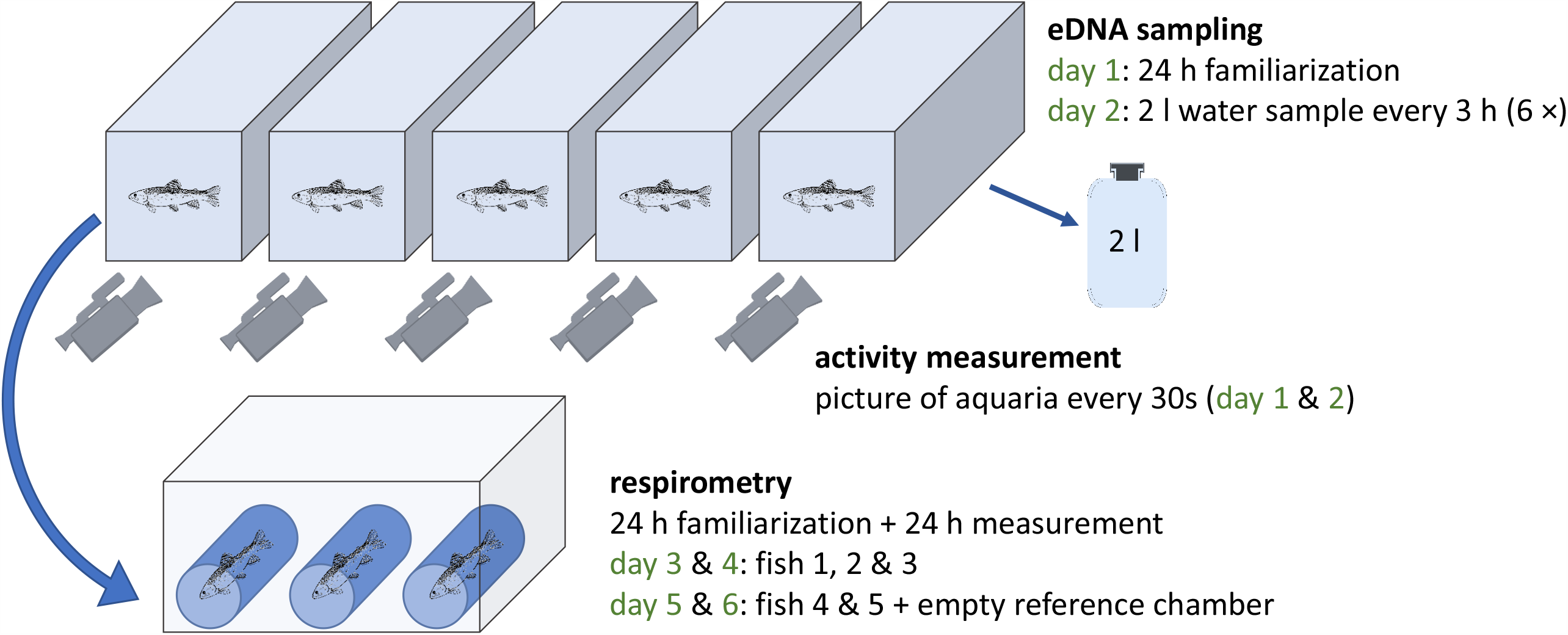
The setup of the aquarium experiment carried out with seven fish species: five individual fish were put in fish tanks for water sampling (eDNA) and activity recordings (day 1 and 2) followed by respirometer measurements (three individuals on days 3 and 4; two individuals plus empty control chamber on days 5 and 6). For *Phoxinus phoxinus* and *Squalius cephalus* the experiment was repeated using groups of three individuals per tank and respirometer chamber.

### 2.3. Water sampling, filtration, and pH

All equipment used for this process was cleaned with sodium hypochlorite (5 %) and rinsed with tap water prior to each use; DNA-free gloves were always worn. On the second day, 2 l water samples were taken every three hours from 9:00 AM to 12:00 AM (six samples) at the back end of each aquarium (opposite to the inflow) using flexible tubes and 2 l wide neck bottles (Fig. 2). Due to the high flow rates (entire water volume replaced every 11 min), the water level in each aquarium self-adjusted automatically after every sampling. The water samples were immediately filtered in an adjacent laboratory using glass microfiber filters (1.2 µm pore width, 47 mm diameter, Whatman GF/C) and one negative control (2 l MilliQ-water) was included per sampling event. Thereafter, the filters were individually placed in 2 ml reaction tubes and stored at –28 °C until further processing in a special diagnostic molecular laboratory at the Department of Zoology, University of Innsbruck (Austria). After each sampling, pH was measured in three arbitrarily selected aquaria using a Hach HQ40 device.

### 2.4. Activity measurement

During the familiarization time (day 1) and between water samplings, fish swimming activity was quantified using a custom-made activity monitoring system consisting of one high-definition USB camera (Ziggi HD Plus, IPEVO.COM) per aquarium. The cameras were placed at the front of each tank and the focus was set towards the back end (Fig. 2). To enable recordings during the night, aquaria were lighted throughout the two recording days. Additionally, white polystyrene plates were used to cover the bottom and the sides to exclude influences from neighboring aquaria and standardize reflections. The signals from the cameras were acquired with a frame rate of 2 frames per minute (fpm) with a macro using the image analysis software FIJI (http://fiji.sc/, a distribution of ImageJ) for MacOS (Schindelin et al., 2012; Rueden et al., 2017). For each aquarium, a region of interest (ROI) excluding the inflow, air-stone and sidewalls was set manually (SM 2). Subsequent frames were arithmetically subtracted and the average gray-scale within the region of interest, as a quantification of fish activity, was extracted from the difference-images. The dataset was manually checked to exclude artifacts produced by changes in illumination (light/dark illumination of the fish), water sampling, measurement of abiotic factors, fogged-up aquarium front and few other camera movements sometimes leading to a changing region of interest in the recordings (SM 2).

### 2.5. Respirometry

A custom-made intermittent-flow respirometer was used (Forstner, 1983; Svendsen et al., 2016) including three measurement chambers placed in a larger tank (Fig. 2). The device was cleaned prior to each fish change using a mixture of 3 % hydrogen peroxide and 3 l of tap water. The volume of each chamber was determined prior to the experiment and oxygen saturation (100 %) and temperature (8 – 9 °C) were kept constant in the tank via an airstone and heating/cooling device (Lauda DLK 10 and Alpha 1, Lauda Germany). The three chambers of the respirometer were connected to the respirometers’ water circuit, constantly pumping O_2_-saturated water from the large tank through the three chambers. For measurements of oxygen consumption, a chamber was cut off from this circuit and a short, closed-loop for this chamber was established. Dissolved oxygen was measured in this chamber every 30 s for a period of 15 min. using a YSI ProODO probe (YSI Inc.) and logged to a computer before the system switched to the next chamber for a 15 min measuring period. On the third day of an experimental run, three of the five fish were placed individually into the chambers avoiding air bubbles and kept there for 24 h for familiarization. On the fourth day, respirometer measurements were carried out for 24 h. Thereafter, the remaining two fish individuals were placed in two measurement chambers for one day of familiarization followed by one day of measurements (days five and six; Fig. 2). The third chamber was left empty, but measured as well, to evaluate potential microorganism-induced oxygen decrease. After the respirometer measurement day, the mass [g] and total length [mm] of each fish were determined before placing them together in a fish tank. For respirometer measurements of fish groups, the three individuals previously sharing an aquarium were put together in a respirometer chamber.

### 2.6. Filter processing and molecular analysis

After defrosting, each filter was soaked with 200 µl of lysis buffer consisting of TES-buffer (0.1 M TRIS, 10 mM EDTA, 2 % sodium dodecyl sulphate; pH 8) and proteinase K (20 mg/ml) in a ratio of 19:1 and incubated at 56 °C over night in a rocking platform. On the next day, filters were transferred with DNA-free forceps to a perforated inset which was repositioned in the top half of the original 2 ml reaction tube and centrifuged for 10 min at 20 000 *g*. Afterwards, filters were discarded and the lysate at the bottom of the reaction tube (300-800 µl) was used for DNA extraction. Insets were cleaned in sodium hypochlorite (2.5 %) for at least 30 min, thoroughly washed with MilliQ-water (10 wash steps) and reused.

DNA extraction was carried out with the Biosprint 96 instrument (Qiagen) using the Biosprint 96 DNA blood Kit (Qiagen) and the Biosprint 96 tissue extraction protocol following the manufacturer’s instructions except for using 100 µl of TE-buffer instead of AE-buffer for DNA elution. Extractions were carried out in 96-well plates and four negative controls (containing TES-buffer instead of lysate) were included per plate. To process the whole lysate volume, a custom DNA-uptake program was set up: three uptake plates were used and 300 µl of lysate, 300 µl AL-buffer and 300 µl isopropanol were mixed per well in each plate. Missing lysate volumes (i.e. if only a total of 400 µl were available after centrifugation) were replaced by TES-buffer. Additionally, 30 µl MagAttract was added per well in the first plate. Using custom “binding” steps of the robotic platform, the DNA contained in the first plate was transferred to the second one. Next, a binding step was carried out in the second plate before transferring and releasing the entire collected DNA into the third plate, which was then used for the Biosprint 96 tissue extraction protocol. After extraction, each eluate was transferred to a 1.5 µl reaction tube for subsequent PCR.

All used primers (Table 1) have been previously published after extensive specificity and sensitivity testing (Thalinger et al., 2016, 2020b) and additional specificity tests were carried out on the digital PCR (dPCR) system (see below) confirming the specificity of the molecular assays under the following conditions: each 22 µl dPCR master mix for droplet generation on the QX200 AutoDG (Biorad) consisted of one-time EvaGreen Supermix (Biorad), 0.25 µM forward and reverse primer (Table 1) and up to 10.5 µl DNA extract. Depending on the results of initial tests with capillary electrophoresis PCR (i.e. the Relative Fluorescence Units (RFU) of the resulting band; see SM 3), extracts were diluted with molecular grade water for dPCR as follows: RFU < 0.2: undiluted; 0.2 ≤ RFU < 1.3: 1:1 dilution; 1.3 ≤ RFU < 2: 1:3 dilution; 2 ≤ RFU: 1:7 dilution. Optimized thermo-cycling conditions were 5 min at 95°C, 40 cycles of 30 s at 95°C, 1 min at 58°C (*O. mykiss, P. phoxinus*, and *S. cephalus*) or 60°C (*C. gobio, S. fontinalis, S. trutta*, and *T. thymallus*), 1 min at 72°C, followed by one step of 5 min at 4°C and 5 min at 90°C. dPCR results were analyzed on the QX200(tm) Droplet Reader with the corresponding QuantaSoftTM Analysis Pro Software (Version 1.7; Biorad). As target signal amplitude varied with the length of the amplified fragment, amplitude thresholds were set individually per primer pair (Table 1) prior to determining target copy numbers per µl for each DNA extract. Per primer pair, a positive (DNA extract from target species) and a negative control (molecular grade water) were included in dPCR, all of which resulted positive and negative, respectively. All filtration and extraction controls resulted negative as well.

**Table 1:**
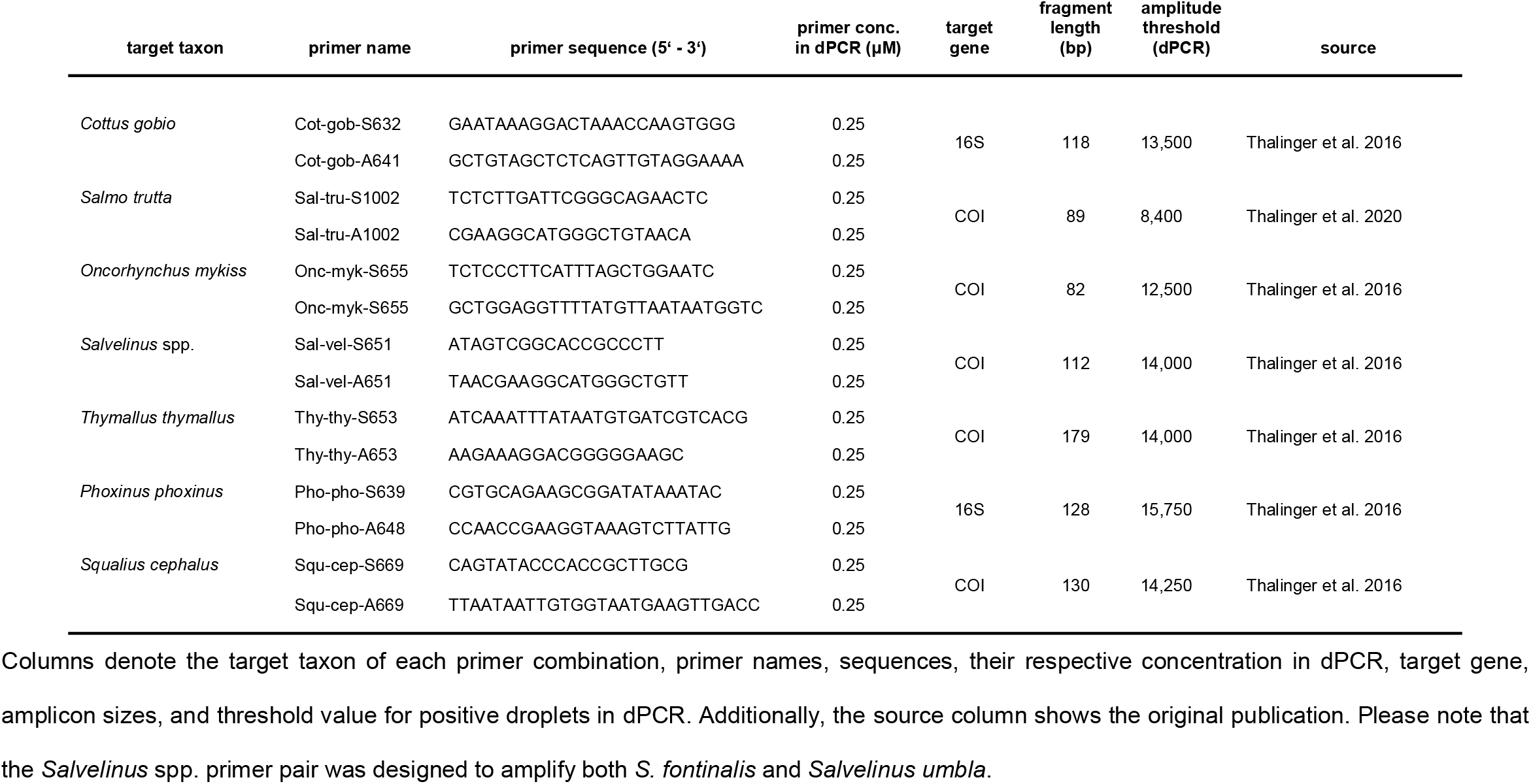
digital PCR assays used to amplify fish eDNA

### 2.7. Statistical analysis

All calculations and visualizations were carried out in R Version 4.0.2 (R Core Team, 2020) using the packages “ggplot2” (Wickham, 2016), “gridExtra” (Auguie, 2017), “ggpubr” (Kassambara, 2019), “lme4” (Bates et al., 2015), “AICcmodavg” (Mazerolle, 2020), “MuMIn” (Barton, 2019), “rsq” (Zhang, 2020) and “sjPlot” (Lüdecke, 2020). As pH was not measured in all aquaria after each water sampling, missing values were estimated by averaging measurements taken at the respective fish tank before and after the skipped time step. If measurements at the first or last water sampling were missing, the values of the following or previous time step, respectively, were carried over.

The cleared activity dataset was visually inspected and summarized for each time step: for example, data obtained during the preceding day were associated with the first eDNA sampling event at 9:00 AM and measurements between 9:00 AM and 12:00 PM were considered relevant for the second water sampling at 12:00 PM. Mean activity was calculated per time interval. No cleared activity data was available for one *S. trutta* and *S. fontinalis* individual, respectively, and for one *P. phoxinus* and *T. thymallus* individual at a single time step each.

The total respirometry dataset was cleared of all 15 min measurement series showing an increase in dissolved O_2_. As this value is expected to decrease linearly over the course of a measurement (Svendsen et al., 2016), a linear regression for the oxygen decrease in a measurement chamber over time was calculated for each measurement series. All intervals for which the obtained values showed an insufficient fit to a linear decrease (R^2^ < 0.8) were also excluded from further analyses. For each of the remaining measurement intervals, oxygen consumption (OC) in mg / h was calculated as *OC* = -*s* × 60 × *vol* where “s” denotes the slope of the linear regression and “vol” the volume of the respective measurement chamber minus the mass of the fish. Per fish species, the obtained value was corrected for the mean oxygen consumption in the empty chamber before calculating total energy use (oxygen consumption × 13.6 J/mg (oxycaloric factor (Brett and Groves, 1979)) per fish. Finally, energy use [J/h] was averaged across the values obtained from individual measurement intervals for each fish and fish group. Due to data clearing, this was not possible for one individual and one group of *C. gobio* and *S. cephalus*, two individuals of *S. fontinalis* and *S. trutta* and three individuals of *T. thymallus*. For these fish, energy use was estimated as the mean of the available values.

Concerning the fish eDNA copy numbers obtained from dPCR, 21 filtered water samples did not lead to an amplification. They were removed from the dataset, as other fish individuals of comparable size and other samplings reliably produced positive results and/or eDNA was detected in celPCR. Hence, processing errors during sampling and in the laboratory were deemed the most likely cause for failing amplification. One group of *P. phoxinus* had to be excluded from further analyses, as two of three individuals were identified as *S. cephalus* when removed from the aquarium after the experiment. To determine whether the pH measurements, mean activity and eDNA copy numbers were significantly influenced by sampling (i.e. time of the day), a one-way repeated measurements ANOVA with rank transformation was calculated for each variable using a combination of fish species and aquarium as random factor. A significant trend could not be detected (Table 2). Despite efforts to standardize the mass of the chosen fish individuals within and between species, fish mass was identified as confounding variable (SM 4). Hence, eDNA copies, mean activity, and energy use were normalized by the mass of the respective fish individual prior to all further analyses.

**Table 2:**
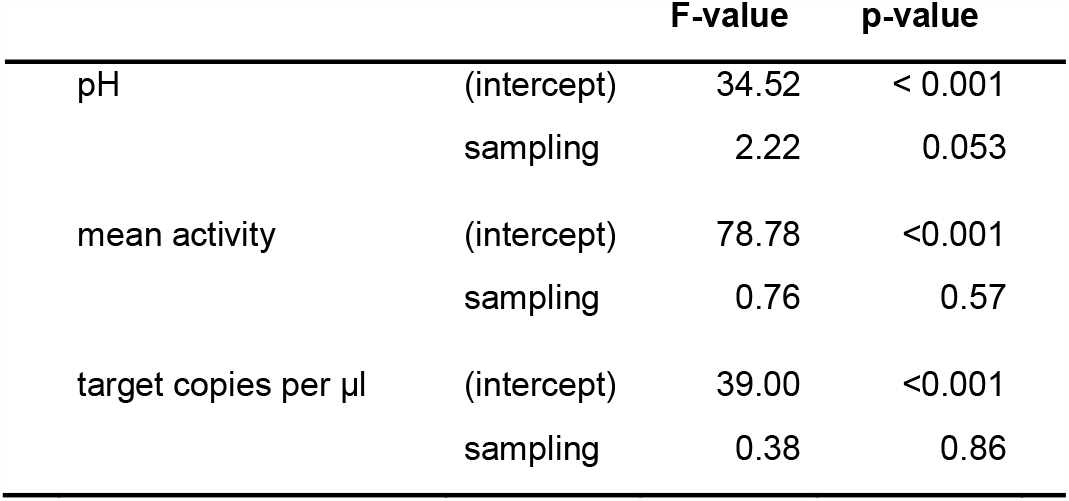
The results of one-way repeated measurements ANOVA with rank transformation examining a potential effect of sampling on pH, mean activity, and target eDNA copy numbers.

|  |  | F-value | p-value |
| --- | --- | --- | --- |
| pH | (intercept) | 34.52 | < 0.001 |
|  | sampling | 2.22 | 0.053 |
| mean activity | (intercept) | 78.78 | <0.001 |
|  | sampling | 0.76 | 0.57 |
| target copies per µl | (intercept) | 39.00 | <0.001 |
|  | sampling | 0.38 | 0.86 |

Generalized Linear Mixed-Effects Models (GLMM) for a Gamma-distributed dependent variable (i.e. eDNA copies) were set up with a log-link function to investigate the effects of mean activity, energy use, fish species, and pH (Faraway, 2016). Data obtained from fish groups were excluded from the comparison of model performance. Fish individuals were used as random intercept to account for repeated measurements and models were fit using Gauss-Hermite quadrature (nAGQ = 20) and the BOBYQA algorithm (Bolker et al., 2009; Powell, 2009). The variable “fish species” was entered via dummy coding into the models using *C. gobio* as base category. Corresponding with the focus of this study to investigate the effect of species identity, energy use, and activity on eDNA shedding, a set of six candidate models was chosen (Table 3). AICc, ΔAICc, and AICc weights (ω) were used to evaluate the strength of the six models for describing the data including marginal and conditional pseudo-R^2^ values (Burnham and Anderson, 2002; Nakagawa and Schielzeth, 2013). Simulated, scaled residuals were calculated based on the best-performing candidate model, (package: DHARMa (Hartig, 2020); function: “simulateResiduals”; n = 1000). The best performing model passed the consecutive check for outliers and overdispersion; the 95%CI for its fixed effects were derived via bootstrapping (200 simulations).

**Table 3:** Covariate structures of the candidate Gamma GLMM with a log link function.

| model # | covariate structure (fixed effects) |
| --- | --- |
| 1 | mean activity + energy use + fish species + pH + sampling |
| 2 | mean activity + energy use + fish species + sampling |
| 3 | mean activity + energy use + fish species |
| 4 | mean activity + energy use |
| 5 | mean activity + fish species |
| 6 | fish species |

The models were compared for their potential to explain the target eDNA copy numbers per gram fish and μl extract obtained from single-fish aquaria with the following parameters: fish species identity (seven species), mean activity (per gram fish), energy use (per gram fish) and pH. In all models, individual fish were included as random effect (random intercept) to account for repeated measurements and the sampling event was included primarily to show its insignificance.

To test the differences between single and grouped fish in the different stages of the experiment, a data subset containing only values obtained from single and grouped *P. phoxinus* and *S. cephalus* was analyzed. Target eDNA copies, energy use, and mean activity (all normalized by fish mass) of the four distinct fish categories were tested for normality and homogeneity of variance with Shapiro-Wilk and Bartlett tests. Then, differences between groups were examined via Kruskal-Wallis tests followed by Wilcoxon Rank Sum tests with Benjamini-Hochberg-corrected p-values. In a final step, target eDNA copies for groups of *P. phoxinus* and *S. cephalus* were predicted using the model previously established for single fish (only possible when not incorporating the random effect of fish individual). Pairwise Wilcoxon tests were used to verify whether there was a significant difference between predicted and measured target eDNA copy numbers for both species separately and combined.

## 3. Results

The mean mass of individually housed fish was 3.06 g ± 1.56 g (SD) and *C. gobio* individuals had the highest mass (5 g ± 2.1 g (SD); Table 4). Water samples from *P. phoxinus* and *T. thymallus* aquaria had the highest eDNA copy numbers per µl extract and gram fish mass (31.13 ± 53.23 (SD) and 47.68 ± 41.13 (SD), respectively; Fig. 3, Table 4). The normalized mean activity was highest for *S. fontinalis* (1.08 ± 0.33 (SD)) and lowest for *C. gobio* (0.34 ± 0.10 (SD); Fig. 3, Table 4). The energy use per gram fish mass was highest for *O. mykiss* (1.81 J/h ± 0.91 J/h (SD)), while *S. fontinalis* and *S. trutta* aquaria had the lowest pH.

**Table 4:**
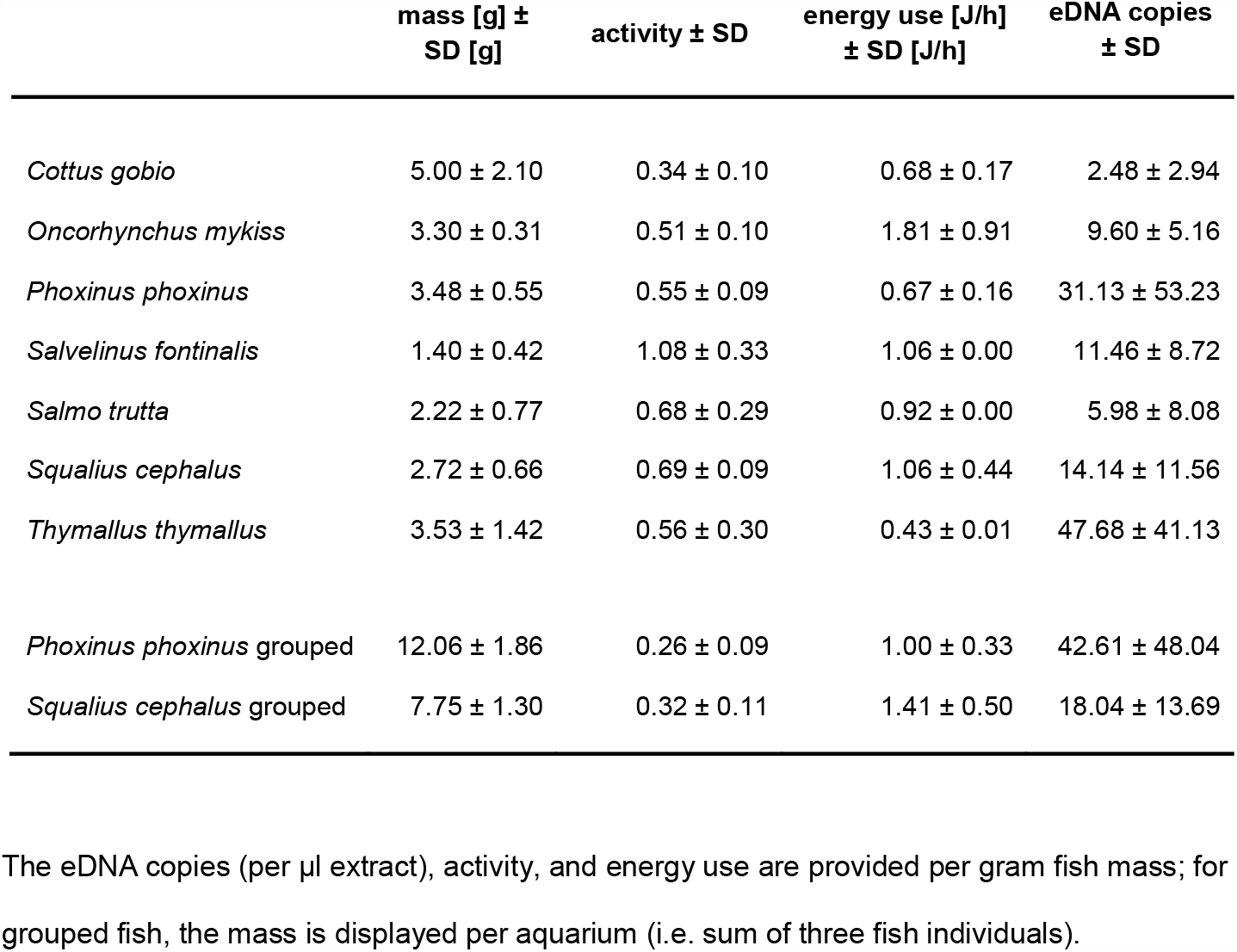
The means and standard deviations of mass, activity, and energy use for each fish species in the experiment.

|  | mass [g] ±<br>SD [g] | activity ± SD | energy use [J/h]<br>± SD [J/h] | eDNA copies<br>± SD |
| --- | --- | --- | --- | --- |
| *Cottus gobio* | 5.00 ± 2.10 | 0.34 ± 0.10 | 0.68 ± 0.17 | 2.48 ± 2.94 |
| *Oncorhynchus mykiss* | 3.30 ± 0.31 | 0.51 ± 0.10 | 1.81 ± 0.91 | 9.60 ± 5.16 |
| *Phoxinus phoxinus* | 3.48 ± 0.55 | 0.55 ± 0.09 | 0.67 ± 0.16 | 31.13 ± 53.23 |
| *Salvelinus fontinalis* | 1.40 ± 0.42 | 1.08 ± 0.33 | 1.06 ± 0.00 | 11.46 ± 8.72 |
| *Salmo trutta* | 2.22 ± 0.77 | 0.68 ± 0.29 | 0.92 ± 0.00 | 5.98 ± 8.08 |
| *Squalius cephalus* | 2.72 ± 0.66 | 0.69 ± 0.09 | 1.06 ± 0.44 | 14.14 ± 11.56 |
| *Thymallus thymallus* | 3.53 ± 1.42 | 0.56 ± 0.30 | 0.43 ± 0.01 | 47.68 ± 41.13 |
| *Phoxinus phoxinus* grouped | 12.06 ± 1.86 | 0.26 ± 0.09 | 1.00 ± 0.33 | 42.61 ± 48.04 |
| *Squalius cephalus* grouped | 7.75 ± 1.30 | 0.32 ± 0.11 | 1.41 ± 0.50 | 18.04 ± 13.69 |

The eDNA copies (per µl extract), activity, and energy use are provided per gram fish mass; for grouped fish, the mass is displayed per aquarium (i.e. sum of three fish individuals).

**Figure 3:**
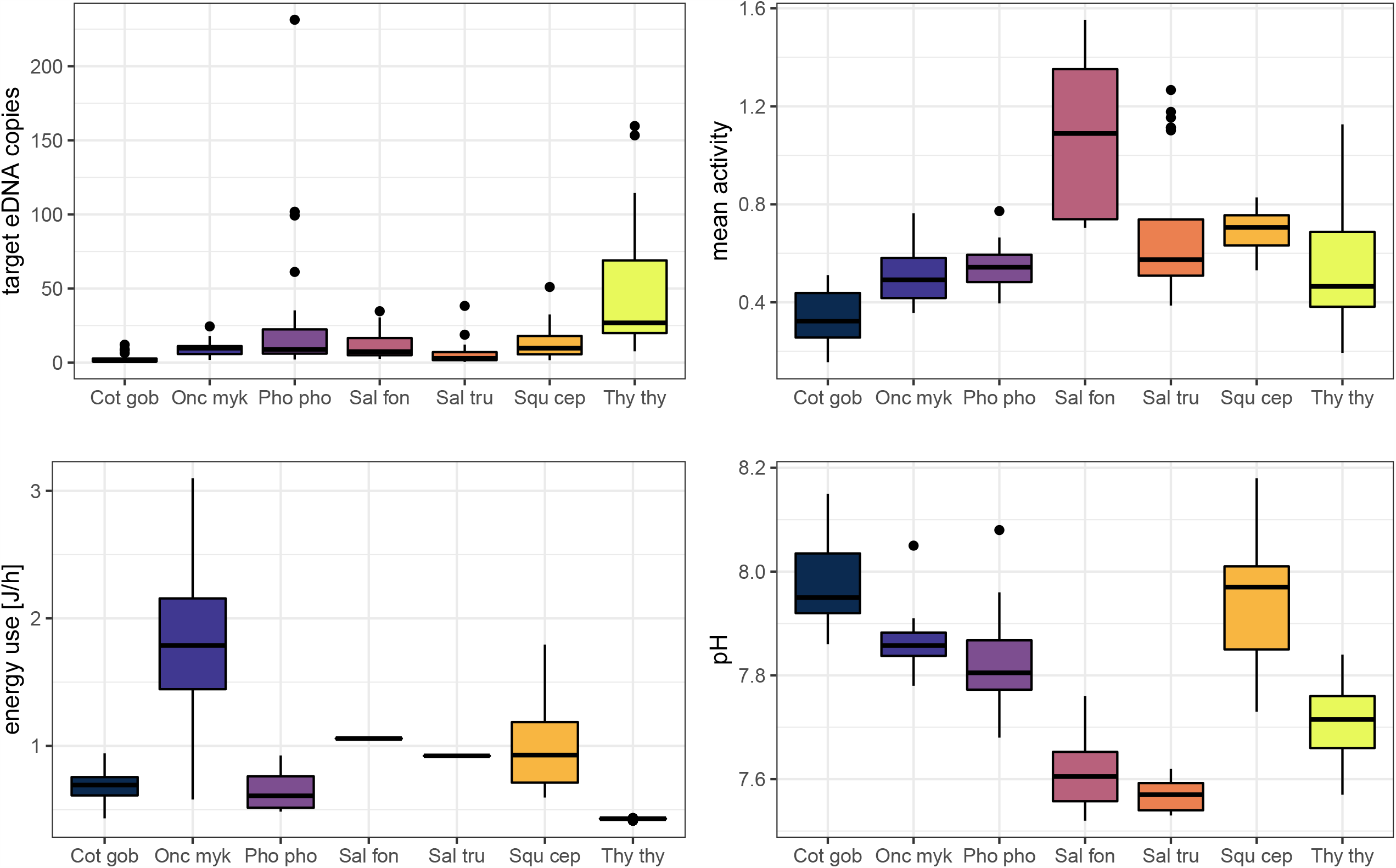
Key parameters obtained during the experiment for single fish. Boxplots display target eDNA copies per µl extract, energy use [J/h], mean activity, and pH per fish species. Fish species are abbreviated: “Cot gob”: *Cottus gobio*; “Onc myk”: *Oncorhynchus mykiss*; “Pho pho”: *Phoxinus phoxinus*; “Sal fon”: *Salvelinus fontinalis*; “Sal tru”: *Salmo trutta*; “Squ cep”: *Squalius cephalus*; “Thy thy”: *Thymallus thymallus*. The variables target eDNA copies, mean activity, and energy use were normalized by fish mass to control for the effect of this confounding variable.

The ΔAICc-based comparison of model weight (single fish only) resulted in model #3 outperforming five other candidate models (Table 3 and Table 5). Therein, mean activity, energy use, and fish species were contained as explanatory variables (conditional pseudo-R^2^ = 0.59). Increased activity had a significantly positive effect on eDNA copy numbers (p < 0.05) and *P. phoxinus, S. cephalus*, and *T. thymallus* displayed significantly higher copy numbers compared to *C. gobio* (base group) after controlling for the effect of fish mass. The relationship between energy use and copy numbers was also positive, but not significant (p = 0.08; Table 6 and Fig. 4).

**Table 5:**
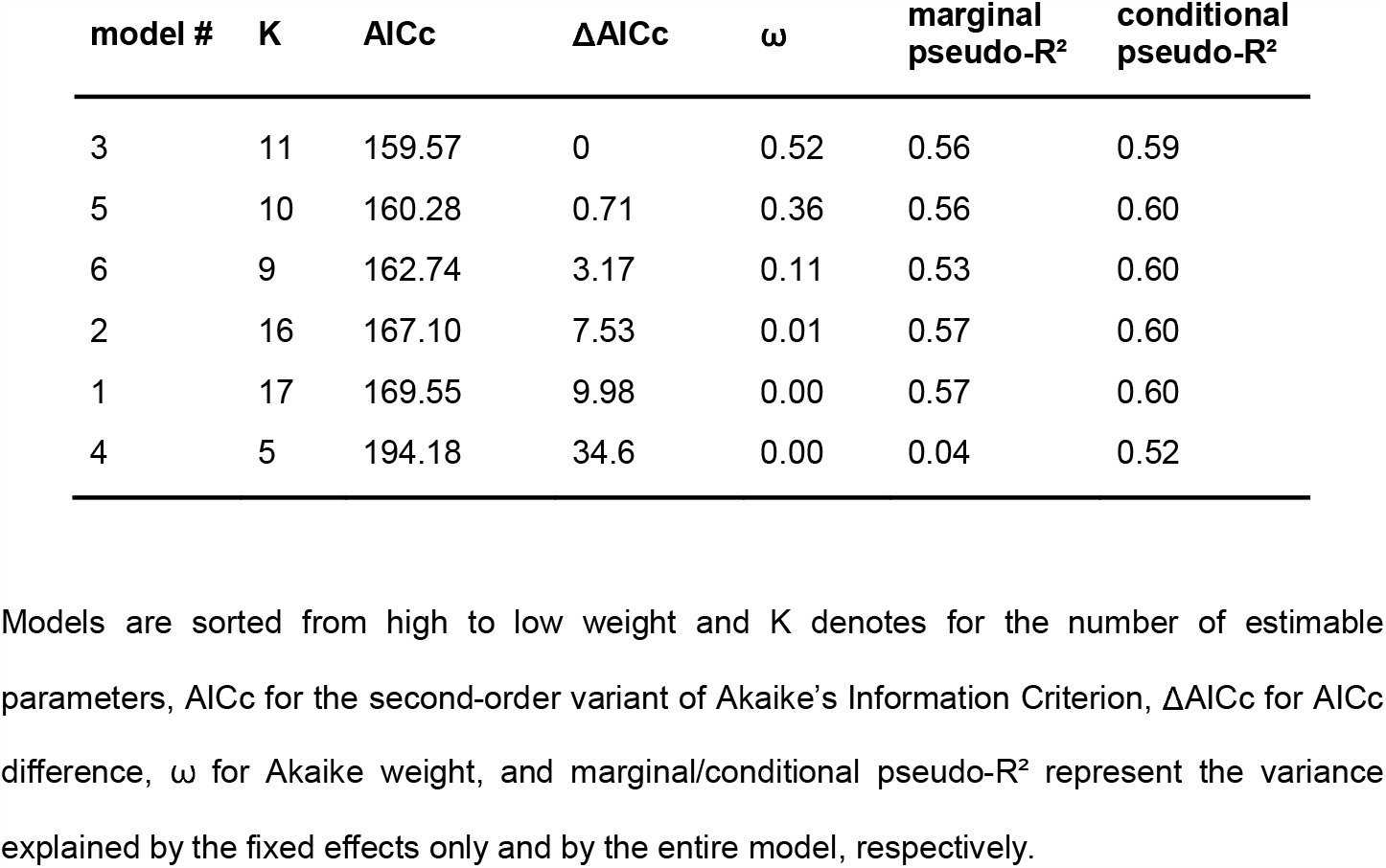
Results of the ordinal ranking based on ΔAICc for the GLMM (Table 3).

| model # | K | AICc | $\Delta AICc$ | $\omega$ | marginal pseudo-R<sup>2</sup> | conditional pseudo-R<sup>2</sup> |
| --- | --- | --- | --- | --- | --- | --- |
| 3 | 11 | 159.57 | 0 | 0.52 | 0.56 | 0.59 |
| 5 | 10 | 160.28 | 0.71 | 0.36 | 0.56 | 0.60 |
| 6 | 9 | 162.74 | 3.17 | 0.11 | 0.53 | 0.60 |
| 2 | 16 | 167.10 | 7.53 | 0.01 | 0.57 | 0.60 |
| 1 | 17 | 169.55 | 9.98 | 0.00 | 0.57 | 0.60 |
| 4 | 5 | 194.18 | 34.6 | 0.00 | 0.04 | 0.52 |

Models are sorted from high to low weight and K denotes for the number of estimable parameters, AICc for the second-order variant of Akaike's Information Criterion, $\Delta AICc$ for AICc difference, $\omega$ for Akaike weight, and marginal/conditional pseudo-R<sup>2</sup> represent the variance explained by the fixed effects only and by the entire model, respectively.

**Table 6:** The highest weight (ω = 0.52) GLMM (model #3) describing the measured eDNA copy numbers via A) the fixed effects: mean activity, energy use and fish species identity, and B) the random effect fish individual (31 groups, σ = 0.88).

| A) | parameter estimate | standard error | lower 95% CI | upper 95% CI | t-value | p-value |
| --- | --- | --- | --- | --- | --- | --- |
| intercept | 0.19 | 0.31 | -0.55 | 0.85 | 0.63 | 0.53 |
| mean activity | 1.00 | 0.42 | 0.19 | 1.95 | 2.39 | 0.02* |
| energy use [J/h] | 0.41 | 0.23 | -0.07 | 0.94 | 1.77 | 0.08 |
| *Oncorhynchus mykiss* | 0.79 | 0.43 | -0.21 | 1.57 | 1.85 | 0.06 |
| *Phoxinus phoxinus* | 2.40 | 0.35 | 1.71 | 3.17 | 6.78 | < 0.001*** |
| *Salvelinus fontinalis* | 0.65 | 0.47 | -0.45 | 1.67 | 1.38 | 0.17 |
| *Salmo trutta* | 0.32 | 0.38 | -0.58 | 1.28 | 0.83 | 0.40 |
| *Squalius cephalus* | 1.25 | 0.38 | 0.41 | 2.01 | 3.28 | < 0.01** |
| *Thymallus thymallus* | 2.93 | 0.33 | 2.30 | 3.58 | 8.75 | < 0.001*** |
| B) | variance | standard deviation |  |  |  |  |
| fish individual (intercept) | 0.06 | 0.24 |  |  |  |  |

Significant p-values of fish species in the model refer to a significant difference between *Cottus gobio* (used as base category for dummy coding) and the respective fish species.

**Figure 4:**
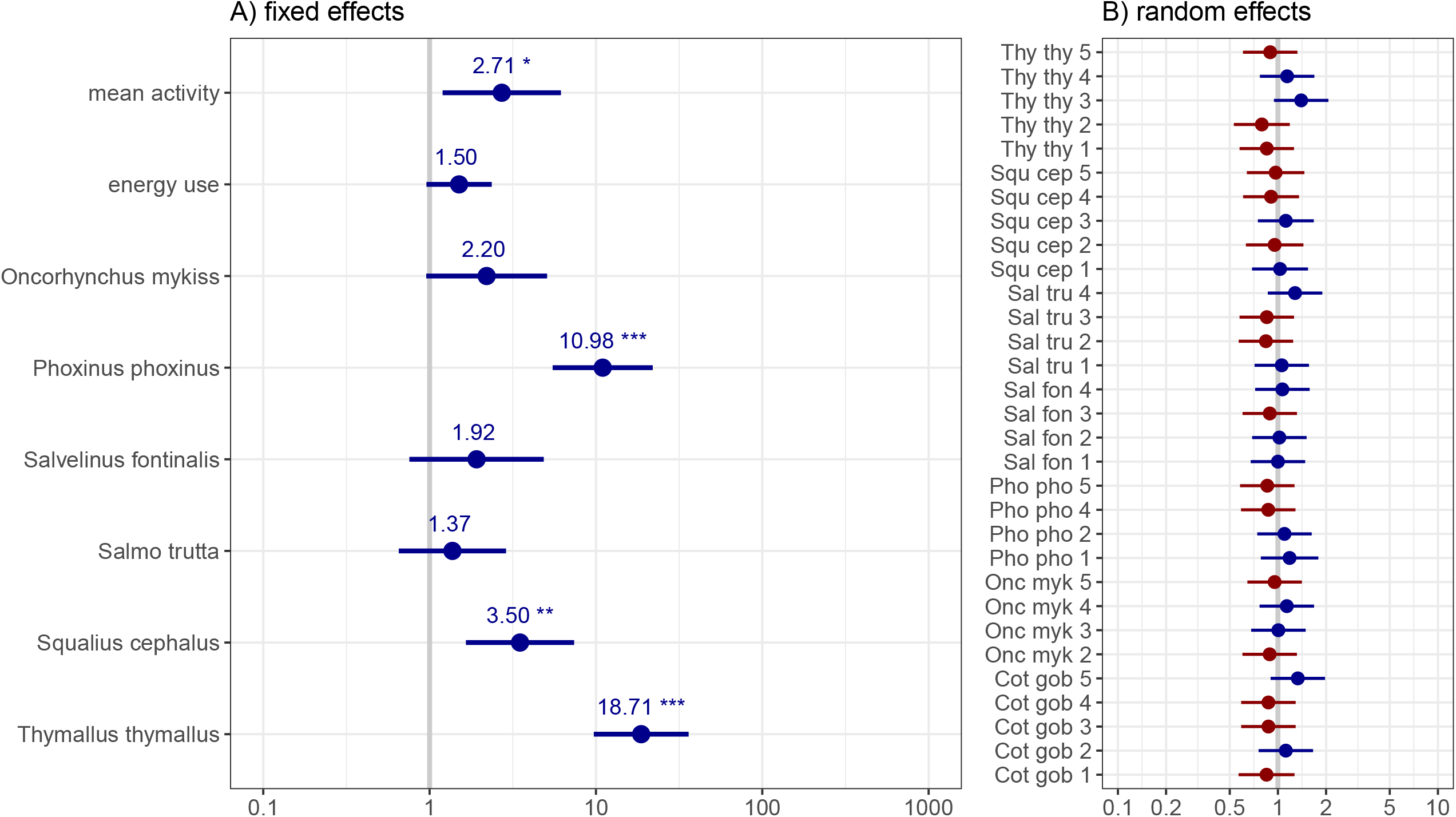
Graphic representation of the GLMM estimates (model #3) best describing the obtained target eDNA copy numbers: A) fixed effects and B) random effect of individual fish. Coefficients are exponentiated, significance codes of denoted fish species indicate differences in comparison to the base category *Cottus gobio*, whiskers display the 95%-CI. Fish species are abbreviated: “Cot gob”: *Cottus gobio*; “Onc myk”: *Oncorhynchus mykiss*; “Pho pho”: *Phoxinus phoxinus*; “Sal fon”: *Salvelinus fontinalis*; “Sal tru”: *Salmo trutta*; “Squ cep”: *Squalius cephalus*; “Thy thy”: *Thymallus thymallus* in addition to individual numbers from 1 to 5.

For single and grouped individuals of *P. phoxinus* and *S. cephalus*, target eDNA copies per gram fish were significantly higher for grouped fish in general (28.96 ± 35.44 (SD) compared to 22.44 ± 38.64 (SD); Chi^2^ = 5.96; p < 0.05). Specifically, they were significantly higher for grouped *P. phoxinus* (42.61 ± 48.04 (SD)) compared to single *P. phoxinus* and single and grouped *S. cephalus* and characterized by few outliers with particularly high eDNA concentration (Fig. 3 and Fig. 5). Significant differences were also detected between the four groups regarding mean activity (Chi^2^ = 80.95; p < 0.001) and energy use (Chi^2^ = 36.77; p < 0.001): mean activity was significantly higher when fish were kept solitary compared to having them in groups for both species (p < 0.001). Contrastingly, energy use was significantly higher for grouped individuals of *P. phoxinus* and *S. cephalus* (p < 0.01).

**Figure 5:**
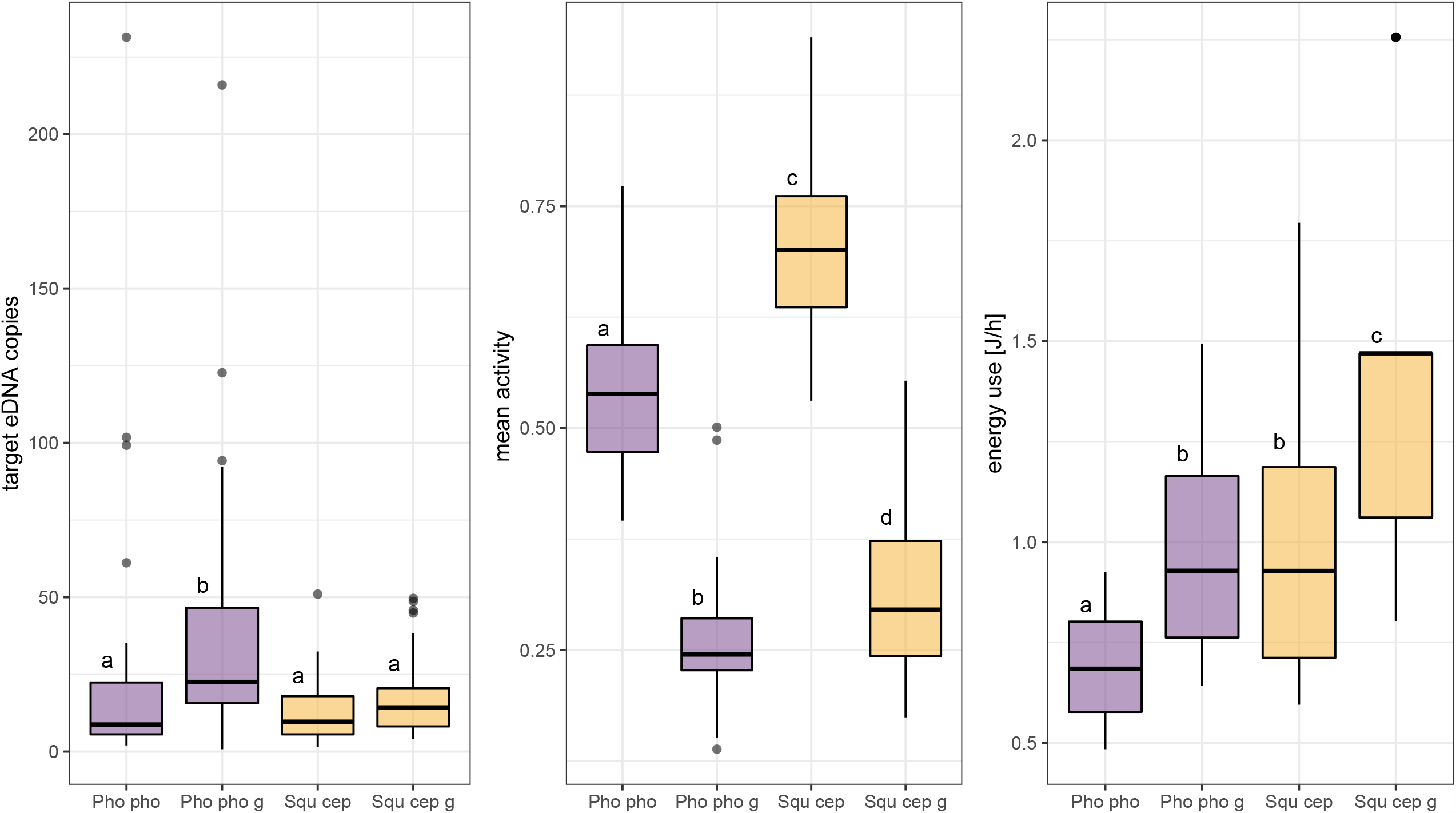
Comparison of target eDNA copies, mean activity, and energy use (normalized by fish mass) in aquaria obtained from single and grouped individuals of *Phoxinus phoxinus* and *Squalius cephalus*. Different lower case letters above boxplots code for significant differences (p < 0.05) between categories, which are abbreviated as: “Pho pho”: *Phoxinus phoxinus* (single fish); “Pho pho g”: *Phoxinus phoxinus* grouped fish; “Squ cep”: *Squalius cephalus* (single fish); “Squ cep g”: *Squalius cephalus* grouped fish.

To test the suitability of model #3 for describing eDNA shedding also for grouped fish, model #3-predicted eDNA copies were compared to the measured copy numbers in the group treatments. For the two species combined, there was no significant difference between predicted and measured copy numbers (W = 1612, p = 0.35). For *P. phoxinus* alone, no such difference was detected either (W = 274; p = 0.78; Fig. 6), while predicted and measured copy numbers of *S. cephalus* showed a significant difference (W = 609; p < 0.05; Fig. 6).

**Figure 6:**
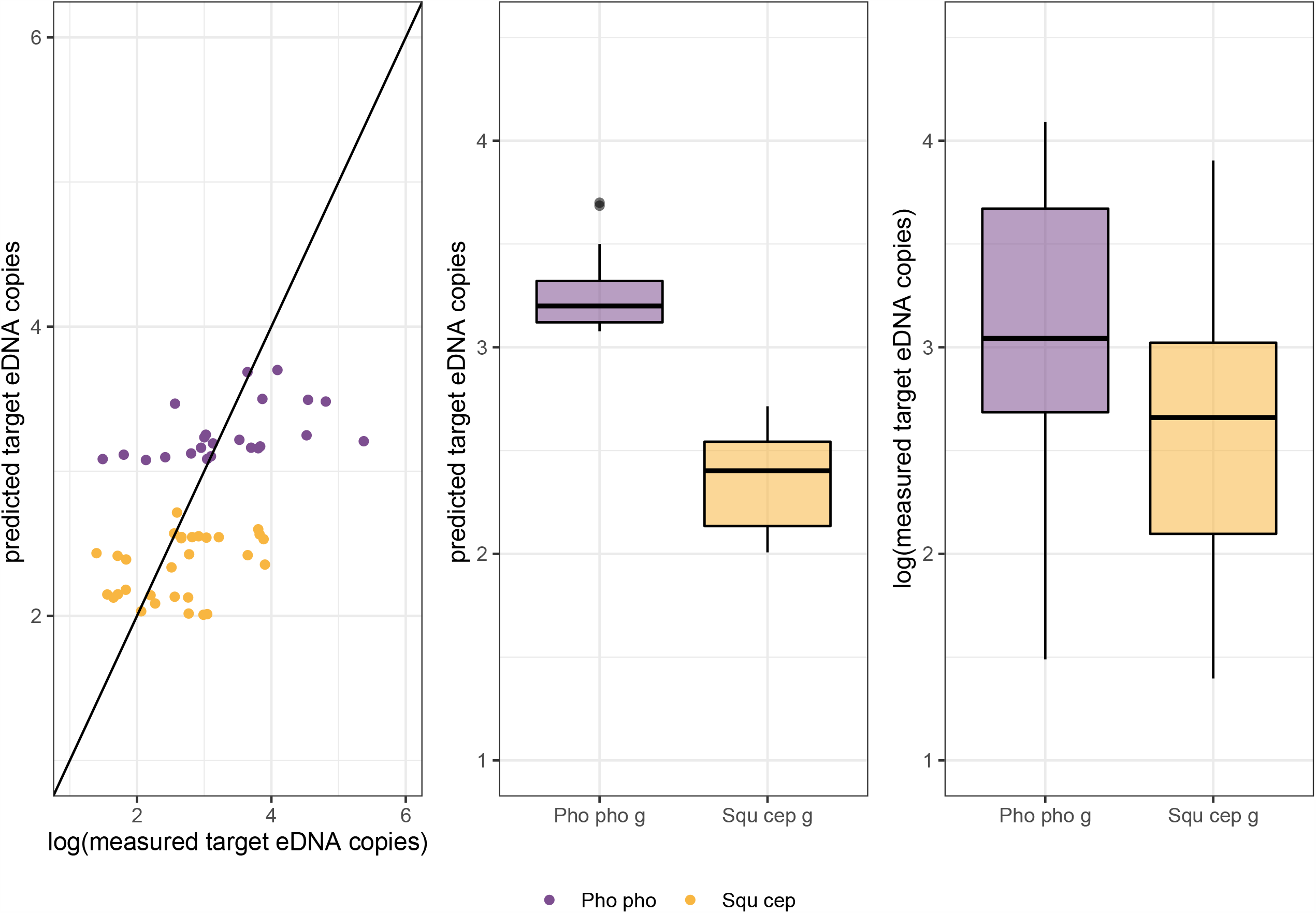
For groups of *Phoxinus phoxinus* (Pho pho g) and *Squalius cephalus* (Squ cep g) measured and predicted copy numbers are plotted: left: against each other; middle: predicted copy numbers are compared between species; right: comparison of measured copy numbers between the two species. The measured copy numbers were log-transformed to enable a direct comparison with the values predicted by the Gamma GLMM with log-link function; the random effect of individual fish could not be taken into account for this prediction. For *S. cephalus* a significant difference between measured and predicted copy numbers was detected (W = 609; p < 0.05).

## 4. Discussion

This experiment confirms the hypothesized positive relationship between eDNA shedding and fish activity. The species identity and thereby associated physiological differences were found to influence the amount of released eDNA, and the positive relationship between energy use and eDNA signals was not significant. Furthermore, our data show that models of eDNA shedding cannot always be generalized from experiments with individual fish to fish groups. For a conclusive habitat-scale estimation of fish communities with eDNA-based methods it is therefore necessary to incorporate species physiology and behavior into the analysis.

In early aquarium experiments, the strongest eDNA signals were found right after the introduction of fish into tanks without water circulation and often explained by elevated stress levels through handling and adaption to the new environment (Takahara et al., 2012; Maruyama et al., 2014, 2019; Klymus et al., 2015). Hence, many recent studies allow for one or several days of accommodation prior to eDNA sampling (Lacoursière-Roussel et al., 2016; Jo et al., 2019; Takeuchi et al., 2019). We can confirm the positive relationship between fish activity (i.e. movement) and eDNA shedding independent of the introductory phase of an experiment. However, it was not possible to determine the actual reason for the elevated eDNA levels associated with higher activity, as both higher metabolic rates during movement and higher water volumes shearing against the fish body could be responsible for this effect. For eDNA-based field studies, this result indicates that signals emitted by highly active fish (e.g. during spawning or predatory behavior) potentially mimic higher levels of fish biomass.

Energy use in a resting state as measured with an intermittent-flow respirometer, was also positively correlated with eDNA production, albeit not significant. In case this trend is confirmed in the future, it could be attributed to the higher metabolic rate and larger gill size of active species in combination with higher water volumes pumped through them (Wegner et al., 2009). However, the elevated eDNA signals could also stem from other physiological processes (e.g. defecation), which are known to positively influence eDNA production rates (Klymus et al., 2015). As fish were not fed during the entire experiment, the latter factor is potentially negligible, albeit it might substantially influence eDNA levels under natural conditions. Except for *T. thymallus*, the energy use of the species preferring microhabitats with strong currents and preying on fish as adults (primarily *O. mykiss* and *S. cephalus*) was higher than for *C. gobio* and *P. phoxinus*. This is in concordance with general differences in resting metabolic rates between these ecological guilds (Roberts, 1975; Johnston et al., 1988; Killen et al., 2010). In this experiment, the smaller sized *S. fontinalis* and *S. trutta* individuals, were more difficult to measure with the chosen respirometer setup (i.e. fewer measurements passed our quality filtering), which could be the cause for the weak relationship between energy use and eDNA copy numbers. In the future, more emphasis should be placed on a ratio of 20-50 between the volume of the measurement chamber in the respirometer and the fish individual to facilitate respirometer measurements (Svendsen et al., 2016).

There were distinct differences in eDNA shedding between the species, with *T. thymallus, P. phoxinus*, and *S. cephalus* emitting the most eDNA. The adaptation to habitats with stronger currents (Freyhof and Kottelat, 2007), namely an increased mucus production in combination with comparably large scales, might explain this result for *T. thymallus* and *S. cephalus*. The underlying taxonomy could also contribute to this pattern if cyprinids (*P. phoxinus* and *S. cephalus*) generally release more DNA into the surrounding water via their gills, feces or mucus. Another explanation for the high eDNA shedding of cyprinids in this experiment could be the stress induced by solitary housing. The model estimating eDNA concentrations for individual fish could not fully explain the findings obtained for grouped fish: the activity of both *P. phoxinus* and *S. cephalus* was significantly lower when fish were held in groups, while their energy use was significantly higher. A change in measurement precision regarding activity and energy use (respirometer more precise, activity measurement less precise for fish groups) could explain these contradictory results. Nevertheless, eDNA copies of grouped *P. phoxinus* individuals did not differ from values predicted with a model based on single fish.

Generally, the measured eDNA concentrations per µl DNA extract were right-skewed and a few exceptionally high values showed a considerable influence on the size of standard deviations. These results were independent of fish handling and stress during the introduction phase as eDNA sampling started only after 24 h and the aquaria had constant flow with the entire volume being renewed every 11 min. Such “outliers” were also detected in other aquarium experiments (Klymus et al., 2015; Wilcox et al., 2016) and cell-conglomerates released into the surrounding water were previously deemed responsible for this pattern (Wilcox et al., 2016). Additionally, the size distribution of eDNA particles (starting from < 0.2 µm and exceeding >180 µm) and commonly detected fragment sizes suggest intact cells or organelles as the primary source of eDNA in the water column (reviewed by Harrison et al., 2019). Our data support the hypothesis of constant eDNA shedding rates at constant environmental conditions indicating the potential to determine this variable for a broad range of species and add to the interpretation of field sampling results. We could not observe any effects of sampling time, possibly due to the constant illumination of the aquaria. Hence, this aspect is not necessarily transferable to natural environments where fish are known to exhibit distinct diurnal movement patterns (Helfman, 1986).

The influence of fish mass on eDNA concentrations was not in the focus of this experiment and fish individuals were as similar in size/mass as possible. However, adult fish of *P. phoxinus* and *C. gobio* are considerably smaller in comparison to the other species (Freyhof and Kottelat, 2007) and the respective juveniles were thus closer to sexual maturity. Hence, the allometric change of metabolic processes (Brown et al., 2004) could be an alternative explanation for the comparably low energy use of these two species. For studies investigating eDNA shedding directly from live animals, biomass will always be an influential and potentially confounding variable and should thus be considered carefully already during experimental design. Recently, the allometrically scaled mass was found to be the best index variable for describing eDNA concentrations in lakes (Yates et al., 2020); since excretion rate, metabolic rate and surface area all scale allometrically too (Brown et al., 2004; O’Shea et al., 2006; Vanni and McIntyre, 2016), future experiments could greatly benefit from the incorporation of this concept. For activity measurements via videotaping, fish length had to be used as an index variable. In this context considerations of body shapes and fins, which differ a lot between taxa (Freyhof and Kottelat, 2007), are also advisable. Finally, individual differences are well documented for fish behavior and metabolic rates (Metcalfe et al., 2016). The number or study animals in future experiments should thus be increased to better control for such effects within a species.

Our results demonstrate that for the successful application of eDNA-based methods on a habitat scale it is necessary to incorporate fish physiology and behavior not only in the study design and sampling process (e.g. by sampling at different depths and in different micro-habitats (Littlefair et al., 2020)), but also during data analysis (Barnes and Turner, 2016; Thalinger et al., 2020a). Seasonal patterns could have a much stronger effect on eDNA concentrations in the water column as previously assumed: for instance, many cyprinids in European freshwaters seek calm areas without current during the winter. Their eDNA is less likely to spread through the water column and additionally, their decreased activity lowers the detection probability even further. In the future, the eDNA shedding of diverse fish species and families in relation to their biomass, activity, and energy use should be investigated to deepen our understanding of taxon-specific effects. Until then, estimations of fish biomass from eDNA quantities in field-collected samples should at least take distinct physiology and behavior into account, especially for comparative analyses between species or seasons.

## 5 Acknowledgements

This research was conducted within the eDNA-Alpfish project funded by the Austria Research Promotion Agency (FFG); project number 853219. We thank R. Vogt for his support during the experiment, M. Böcker for assistance with the background literature, J. Harvie for input on the statistic analysis, and C. Moritz and D. Kirschner for their help in obtaining the *C. gobio* individuals. We thank two anonymous reviewers for their extensive, constructive feedback on the original manuscript. This manuscript has been uploaded as preprint to bioRxiv https://doi.org/10.1101/2020.10.28.359653 (Thalinger et al., 2020c).

## 7. Author contribution statement

MT and JW, conceived the study; the experiment was designed by BT, MT, TS and JW, and carried out by AR and AT under the supervision of BT and JW. TS was responsible for the processing of activity data; AR, AT and YP were responsible for laboratory processing of the eDNA samples under the supervision of BT who also carried out statistical analysis and wrote the first draft of the manuscript which was revised by all co-authors.

## 8. Data Availability Statement

All data on eDNA signals, fish activity, energy use, fish mass and pH have been uploaded to Figshare and are available at https://doi.org/10.6084/m9.figshare.13151180.v1

## References

Ángeles Esteban, M. (2012). An Overview of the Immunological Defenses in Fish Skin. ISRN Immunol. 2012, 1–29. doi:10.5402/2012/853470.

Auguie, B. (2017). gridExtra: Miscellaneous Functions for “Grid” Graphics. Available at: https://cran.r-project.org/package=gridExtra.

Barnes, M. A., and Turner, C. R. (2016). The ecology of environmental DNA and implications for conservation genetics. Conserv. Genet. 17, 1–17. doi:10.1007/s10592-015-0775-4.

Barton, K. (2019). MuMIn: Multi-Model Inference. Available at: https://cran.r-project.org/package=MuMIn.

Bates, D., Mächler, M., Bolker, B., and Walker, S. (2015). Fitting Linear Mixed-Effects Models Using lme4. J. Stat. Softw. 67, 1--48. doi:10.18637/jss.v067.i01.

Bolker, B. M., Brooks, M. E., Clark, C. J., Geange, S. W., Poulsen, J. R., Stevens, M. H. H., et al. (2009). Generalized linear mixed models: a practical guide for ecology and evolution. Trends Ecol. Evol. 24, 127–135. doi:10.1016/j.tree.2008.10.008.

Brett, J. R., and Groves, T. D. D. (1979). “Physiological energetics,” in Fish Physiology Vol. 8, Bioenergetics and Growt., eds. W. S. Hoar, D. J. Randall, and J. R. Brett (New York: Academic Press). Available at: https://trove.nla.gov.au/work/18117950.

Brown, J. H., Gillooly, J. F., Allen, A. P., Savage, V. M., and West, G. B. (2004). Toward a metabolic theory of ecology. in Ecology (Ecological Society of America), 1771–1789. doi:10.1890/03-9000.

Burnham, K. P., and Anderson, D. R. (2002). Model Selection and Multimodel Inference. 2nd ed. New York: Springer doi:10.1007/b97636.

Bylemans, J., Furlan, E. M., Hardy, C. M., McGuffie, P., Lintermans, M., and Gleeson, D. M. (2017). An environmental DNA-based method for monitoring spawning activity: a case study, using the endangered Macquarie perch (Macquaria australasica). Methods Ecol. Evol. 8, 646–655. doi:10.1111/2041-210X.12709.

Deiner, K., Bik, H. M., Mächler, E., Seymour, M., Lacoursière-Roussel, A., Altermatt, F., et al. (2017). Environmental DNA metabarcoding: Transforming how we survey animal and plant communities. Mol. Ecol. 26, 5872–5895. doi:10.1111/mec.14350.

Evans, N. T., Olds, B. P., Renshaw, M. A., Turner, C. R., Li, Y., Jerde, C. L., et al. (2016). Quantification of mesocosm fish and amphibian species diversity via environmental DNA metabarcoding. Mol. Ecol. Resour. 16, 29–41. doi:10.1111/1755-0998.12433.

Evans, N. T., Shirey, P. D., Wieringa, J. G., Mahon, A. R., and Lamberti, G. A. (2017). Comparative Cost and Effort of Fish Distribution Detection via Environmental DNA Analysis and Electrofishing. Fisheries 42, 90–99. doi:10.1080/03632415.2017.1276329.

Faraway, J. J. (2016). Extending the Linear Model with R. Generalized Linear, Mixed Effects and Nonparametric Regression Models. 2nd ed. New York: Chapman and Hall/CRC doi:10.1201/9781315382722.

Forstner, H. (1983). “An Automated Multiple-Chamber Intermittent-Flow Respirometer,” in Polarographic Oxygen Sensors (Springer Berlin Heidelberg), 111–126. doi:10.1007/978-3-642-81863-9_12.

Freyhof, J., and Kottelat, M. (2007). Handbook of European freshwater fishes. Springer Science and Business Media LLC doi:10.1007/s10228-007-0012-3.

Goldberg, C. S., Turner, C. R., Deiner, K., Klymus, K. E., Thomsen, P. F., Murphy, M. A., et al. (2016). Critical considerations for the application of environmental DNA methods to detect aquatic species. Methods Ecol. Evol. 7, 1299–1307. doi:10.1111/2041-210X.12595.

Harrison, J. B., Sunday, J. M., and Rogers, S. M. (2019). Predicting the fate of eDNA in the environment and implications for studying biodiversity. Proc. R. Soc. B Biol. Sci. 286. doi:10.1098/rspb.2019.1409.

Hartig, F. (2020). DHARMa: Residual Diagnostics for Hierarchical (Multi-Level / Mixed) Regression Models. Available at: https://cran.r-project.org/package=DHARMa.

Helfman, G. S. (1986). “Fish Behaviour by Day, Night and Twilight,” in The Behaviour of Teleost Fishes (Springer US), 366–387. doi:10.1007/978-1-4684-8261-4_14.

Horiuchi, T., Masuda, R., Murakami, H., Yamamoto, S., and Minamoto, T. (2019). BiomassDdependent emission of environmental DNA in jack mackerel Trachurus japonicus juveniles. J. Fish Biol. 95, jfb.14095. doi:10.1111/jfb.14095.

Huerlimann, R., Cooper, M. K., Edmunds, R. C., Villacorta-Rath, C., Le Port, A., Robson, H. L. A., et al. (2020). Enhancing tropical conservation and ecology research with aquatic environmental DNA methods: an introduction for non-environmental DNA specialists. Anim. Conserv. doi:10.1111/acv.12583.

Jo, T., Arimoto, M., Murakami, H., Masuda, R., and Minamoto, T. (2020). Estimating shedding and decay rates of environmental nuclear DNA with relation to water temperature and biomass. Environ. DNA 2, 140–151. doi:10.1002/edn3.51.

Jo, T., Murakami, H., Yamamoto, S., Masuda, R., and Minamoto, T. (2019). Effect of water temperature and fish biomass on environmental DNA shedding, degradation, and size distribution. Ecol. Evol. 9, 1135–1146. doi:10.1002/ece3.4802.

Johnston, I. A., Camm, J. P., and White, M. (1988). Specialisations of swimming muscles in the pelagic antarctic fish Pleuragramma antarcticum. Mar. Biol. 100, 3–12. doi:10.1007/BF00392949.

Kassambara, A. (2019). ggpubr: “ggplot2” Based Publication Ready Plots. Available at: https://cran.r-project.org/package=ggpubr.

Killen, S. S., Atkinson, D., and Glazier, D. S. (2010). The intraspecific scaling of metabolic rate with body mass in fishes depends on lifestyle and temperature. Ecol. Lett. 13, 184–193. doi:10.1111/j.1461-0248.2009.01415.x.

Klymus, K. E., Richter, C. A., Chapman, D. C., and Paukert, C. (2015). Quantification of eDNA shedding rates from invasive bighead carp Hypophthalmichthys nobilis and silver carp Hypophthalmichthys molitrix. Biol. Conserv. 183, 77–84. doi:10.1016/j.biocon.2014.11.020.

Lacoursière-Roussel, A., Rosabal, M., and Bernatchez, L. (2016). Estimating fish abundance and biomass from eDNA concentrations: variability among capture methods and environmental conditions. Mol. Ecol. Resour. 16, 1401–1414. doi:10.1111/1755-0998.12522.

Leese, F., Altermatt, F., Bouchez, A., Ekrem, T., Hering, D., Meissner, K., et al. (2016). DNAqua-Net: Developing new genetic tools for bioassessment and monitoring of aquatic ecosystems in Europe. Res. Ideas Outcomes 2, e11321. doi:10.3897/rio.2.e11321.

Littlefair, J. E., Hrenchuk, L. E., Blanchfield, P. J., Rennie, M. D., and Cristescu, M. E. (2020). Thermal stratification and fish thermal preference explain vertical eDNA distributions in lakes. Mol. Ecol., mec.15623. doi:10.1111/mec.15623.

Lüdecke, D. (2020). sjPlot: Data Visualization for Statistics in Social Science. Available at: https://cran.r-project.org/package=sjPlot.

Maruyama, A., Nakamura, K., Yamanaka, H., Kondoh, M., and Minamoto, T. (2014). The release rate of environmental DNA from juvenile and adult fish. PLoS One 9. doi:10.1371/journal.pone.0114639.

Maruyama, A., Nakamura, K., Yamanaka, H., Kondoh, M., and Minamoto, T. (2019). Correction: The release rate of environmental DNA from juvenile and adult fish (PLoS ONE (2019) 14:2 (e0212145) DOI: 10.1371/journal.pone.0114639). PLoS One 14. doi:10.1371/journal.pone.0212145.

Mazerolle, M. J. (2020). AICcmodavg: Model selection and multimodel inference based on (Q)AIC(c). Available at: https://cran.r-project.org/package=AICcmodavg.

McCollDGausden, E., Weeks, A., Coleman, R., Robinson, K., Song, S., Raadik, T., et al. (2020). MultiDspecies models reveal that eDNA metabarcoding is more sensitive than backpack electrofishing for conducting fish surveys in freshwater streams. Mol. Ecol. doi:10.1111/mec.15644.

Merkes, C. M., McCalla, S. G., Jensen, N. R., Gaikowski, M. P., and Amberg, J. J. (2014). Persistence of DNA in Carcasses, Slime and Avian Feces May Affect Interpretation of Environmental DNA Data. PLoS One 9, e113346. doi:10.1371/journal.pone.0113346.

Metcalfe, N. B., Van Leeuwen, T. E., and Killen, S. S. (2016). Does individual variation in metabolic phenotype predict fish behaviour and performance? J. Fish Biol. 88, 298–321. doi:10.1111/jfb.12699.

Minamoto, T., Miya, M., Sado, T., Seino, S., Doi, H., Kondoh, M., et al. (2020). An illustrated manual for environmental DNA research: Water sampling guidelines and experimental protocols. Environ. DNA, edn3.121. doi:10.1002/edn3.121.

Mizumoto, H., Urabe, H., Kanbe, T., Fukushima, M., and Araki, H. (2018). Establishing an environmental DNA method to detect and estimate the biomass of Sakhalin taimen, a critically endangered Asian salmonid. Limnology 19, 219–227. doi:10.1007/s10201-017-0535-x.

Muus, B. J., and Dahlström, P. (1968). Süßwasserfische. Munich: BLV Verlagsgesellschaft.

Nakagawa, S., and Schielzeth, H. (2013). A general and simple method for obtaining R2 from generalized linear mixed-effects models. Methods Ecol. Evol. 4, 133–142. doi:10.1111/j.2041-210x.2012.00261.x.

O’Shea, B., Mordue-Luntz, A. J., Fryer, R. J., Pert, C. C., and Bricknell, I. R. (2006). Determination of the surface area of a fish. J. Fish Dis. 29, 437–440. doi:10.1111/j.1365-2761.2006.00728.x.

Pilliod, D. S., Laramie, M. B., MacCoy, D., and Maclean, S. (2019). Integration of eDNADBased Biological Monitoring within the U.S. Geological Survey’s National Streamgage Network. JAWRA J. Am. Water Resour. Assoc. 55, 1505–1518. doi:10.1111/1752-1688.12800.

Pont, D., Rocle, M., Valentini, A., Civade, R., Jean, P., Maire, A., et al. (2018). Environmental DNA reveals quantitative patterns of fish biodiversity in large rivers despite its downstream transportation. Sci. Rep. 8. doi:10.1038/s41598-018-28424-8.

Powell, M. J. D. (2009). The BOBYQA algorithm for bound constrained optimization without derivatives. Rep. No. DAMTP 2009/NA06, 1–39. Available at: http://www.damtp.cam.ac.uk/user/na/NA_papers/NA2009_06.pdf.

R Core Team (2020). R: A Language and Environment for Statistical Computing. Available at: https://www.r-project.org/.

Roberts, J. L. (1975). Active branchial and ram gill ventilation in fishes. BIOL.BULL. 148, 85–105. doi:10.2307/1540652.

Rueden, C. T., Schindelin, J., Hiner, M. C., DeZonia, B. E., Walter, A. E., Arena, E. T., et al. (2017). ImageJ2: ImageJ for the next generation of scientific image data. BMC Bioinformatics 18, 1–26. doi:10.1186/s12859-017-1934-z.

Sassoubre, L. M., Yamahara, K. M., Gardner, L. D., Block, B. A., and Boehm, A. B. (2016). Quantification of Environmental DNA (eDNA) Shedding and Decay Rates for Three Marine Fish. Environ. Sci. Technol. 50, 10456–10464. doi:10.1021/acs.est.6b03114.

Schindelin, J., Arganda-Carreras, I., Frise, E., Kaynig, V., Longair, M., Pietzsch, T., et al. (2012). Fiji: An open-source platform for biological-image analysis. Nat. Methods 9, 676–682. doi:10.1038/nmeth.2019.

Shogren, A. J., Tank, J. L., Andruszkiewicz, E., Olds, B., Mahon, A. R., Jerde, C. L., et al. (2017). Controls on eDNA movement in streams: Transport, Retention, and Resuspension. Sci. Rep. 7, 1–11. doi:10.1038/s41598-017-05223-1.

Sigsgaard, E. E., Carl, H., Møller, P. R., and Thomsen, P. F. (2015). Monitoring the near-extinct European weather loach in Denmark based on environmental DNA from water samples. Biol. Conserv. 183, 46–52. doi:10.1016/j.biocon.2014.11.023.

Spindler, T. (1997). Fischfauna in Osterreich. Ökologie – Gefahrdung – Bioindikation – Fischerei – Gesetzgebung. Umweltbund. Vienna, Austria: Umweltbundesamt.

Strickler, K. M., Fremier, A. K., and Goldberg, C. S. (2015). Quantifying effects of UV-B, temperature, and pH on eDNA degradation in aquatic microcosms. Biol. Conserv. 183, 85–92. doi:10.1016/j.biocon.2014.11.038.

Svendsen, M. B. S., Bushnell, P. G., and Steffensen, J. F. (2016). Design and setup of intermittent-flow respirometry system for aquatic organisms. J. Fish Biol. 88, 26–50. doi:10.1111/jfb.12797.

Takahara, T., Minamoto, T., Yamanaka, H., Doi, H., and Kawabata, Z. (2012). Estimation of Fish Biomass Using Environmental DNA. PLoS One 7, e35868. doi:10.1371/journal.pone.0035868.

Takeuchi, A., Iijima, T., Kakuzen, W., Watanabe, S., Yamada, Y., Okamura, A., et al. (2019). Release of eDNA by different life history stages and during spawning activities of laboratory-reared Japanese eels for interpretation of oceanic survey data. Sci. Rep. 9, 1–9. doi:10.1038/s41598-019-42641-9.

Thalinger, B., Deiner, K., Harper, L., Rees, H., Blackman, R., Sint, D., et al. (2020a). A validation scale to determine the readiness of environmental DNA assays for routine species monitoring. bioRxiv. doi:10.1101/2020.04.27.063990.

Thalinger, B., Kirschner, D., Pütz, Y., Moritz, C., Schwarzenberger, R., Wanzenböck, J., et al. (2020b). Lateral and longitudinal fish environmental DNA distribution in dynamic riverine habitats. Environ. DNA 00, 1–14. doi:10.1002/edn3.171.

Thalinger, B., Oehm, J., Mayr, H., Obwexer, A., Zeisler, C., and Traugott, M. (2016). Molecular prey identification in Central European piscivores. Mol. Ecol. Resour. 16, 123–137. doi:10.1111/1755-0998.12436.

Thalinger, B., Rieder, A., Teuffenbach, A., Putz, Y., Schwerte, T., Wanzenboeck, J., et al. (2020c). The effect of activity, energy use, and species identity on environmental DNA shedding in freshwater fish. bioRxiv. doi:10.1101/2020.10.28.359653.

Tsuji, S., Ushio, M., Sakurai, S., Minamoto, T., and Yamanaka, H. (2017). Water temperature-dependent degradation of environmental DNA and its relation to bacterial abundance. PLoS One 12. doi:10.1371/journal.pone.0176608.

Vanni, M. J., and McIntyre, P. B. (2016). Predicting nutrient excretion of aquatic animals with metabolic ecology and ecological stoichiometry: a global synthesis. Ecology 97, 3460–3471. doi:10.1002/ecy.1582.

Wegner, N. C., Sepulveda, C. A., Bull, K. B., and Graham, J. B. (2009). Gill morphometrics in relation to gas transfer and ram ventilation in high-energy demand teleosts: Scombrids and billfishes. J. Morphol. 271, 36–49. doi:10.1002/jmor.10777.

Wickham, H. (2016). ggplot2: Elegant Graphics for Data Analysis. Springer-Verlag New York Available at: https://ggplot2.tidyverse.org.

Wilcox, T. M., McKelvey, K. S., Young, M. K., Sepulveda, A. J., Shepard, B. B., Jane, S. F., et al. (2016). Understanding environmental DNA detection probabilities: A case study using a stream-dwelling char Salvelinus fontinalis. Biol. Conserv. 194, 209–216. doi:10.1016/j.biocon.2015.12.023.

Yates, M. C., Glaser, D. M., Post, J. R., Cristescu, M. E., Fraser, D. J., and Derry, A. M. (2020). The relationship between eDNA particle concentration and organism abundance in nature is strengthened by allometric scaling. Mol. Ecol., mec.15543. doi:10.1111/mec.15543.

Zhang, D. (2020). rsq: R-Squared and Related Measures. Available at: https://cran.r-project.org/package=rsq.

