## Supplementary material for "The effect of activity, energy use, and species identity on environmental DNA shedding of freshwater fish": SM

**Supplementary Material 1: description of preliminary aquarium experiments**

Prior to the experiment described in the associated manuscript, a trial run was carried out with *Oncorhynchus mykiss* and *Thymallus thymallus* individuals. Per fish species, ten aquaria were used and each of them contained five fish individuals with an average mass per aquarium of 253 g ± 49 g (SD) and 55 g ± 7 g (SD) for *O. mykiss* and *T. thymallus*, respectively. The experiment was repeated for three flow rates (3 l/min, 2 l/min and 0.8 l/min) with 2 l water samples taken for subsequent filtration and eDNA analysis at 12 time points during three days (at 0.5, 3.5, 6.5, 9.5, 12.5, 15.5, 21.5, 27.5, 33.5, 39.5, 45.5 and 51.5 h) after placing the fish into the aquaria. Filter processing was similar to the main experiment, however, the obtained filters where shredded with glass beads of different sizes in a Precellys tissue homogenizer (Bertin Technologies) prior to the actual DNA extraction. To estimate the amount of fish eDNA, all samples were subjected to capillary electrophoresis PCR (celPCR) as described in SM 3 (Thalinger et al., 2019, 2020a, 2020b). We deemed this experimental setup unsuitable to investigate the influence of fish species, activity, and energy use for the following reasons:

- The filters showed different levels of dryness prior to the lysis, filter shredding success was not uniform, and both factors lead to differences between lysate volumes. Therefore, the extraction process itself caused strong variations in the DNA concentration of the extracts.
- For all three flow rates the eDNA signals obtained after celPCR were strong, with average relative fluorescence units (RFU) of 2.75 ± 1.01 (SD) and 3.06 ± 1.17 (SD) for *O. mykiss* and *T. thymallus*, respectively. At such DNA concentrations, the results of celPCR were found to be highly variable and potentially not suitable for quantitative analysis (Thalinger et al., 2020b).
- With five fish individuals per tank, the activity analysis via arithmetically subtracted consecutive frames was deemed too error-prone.
- The size of the measurement chambers of the intermittent-flow respirometer was not suitable for five fish individuals at once.
- Stress experienced by non-schooling fish species reared in groups in the aquaria and the respirometer was deemed likely to confound any results.

Therefore, the main experiment was carried out with one fish individual per aquarium and fish groups only for schooling cyprinids. The flow rate was increased to 5.45 l/min, which was the maximum rate possible whilst still providing a constant water temperature of 15°C at the chosen experimental setup. Finally, filter shredding was omitted, and DNA was quantified via digital PCR.

**Supplementary Material 2: Regions of interest (ROI) for activity measurements**

Pictures are sorted per aquarium and ROI are marked in every image. Due to external influences such as slight camera movement or a changing position of the air-stone, ROI were sometimes subjected to slight changes.

**Aquarium 1**

1. *Phoxinus phoxinus* single and group, *Squalius cephalus* single and group, *Oncorhynchus mykiss*

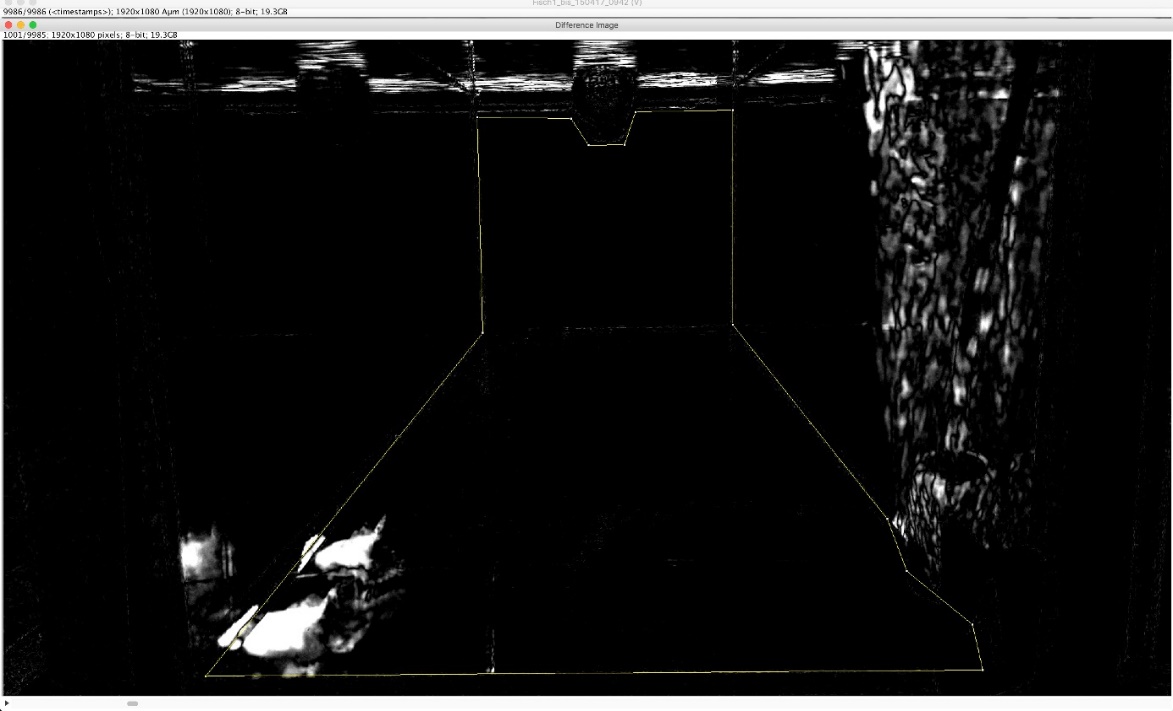

2. *Cottus gobio*, *Salmo trutta*, *Salvelinus fontinalis*

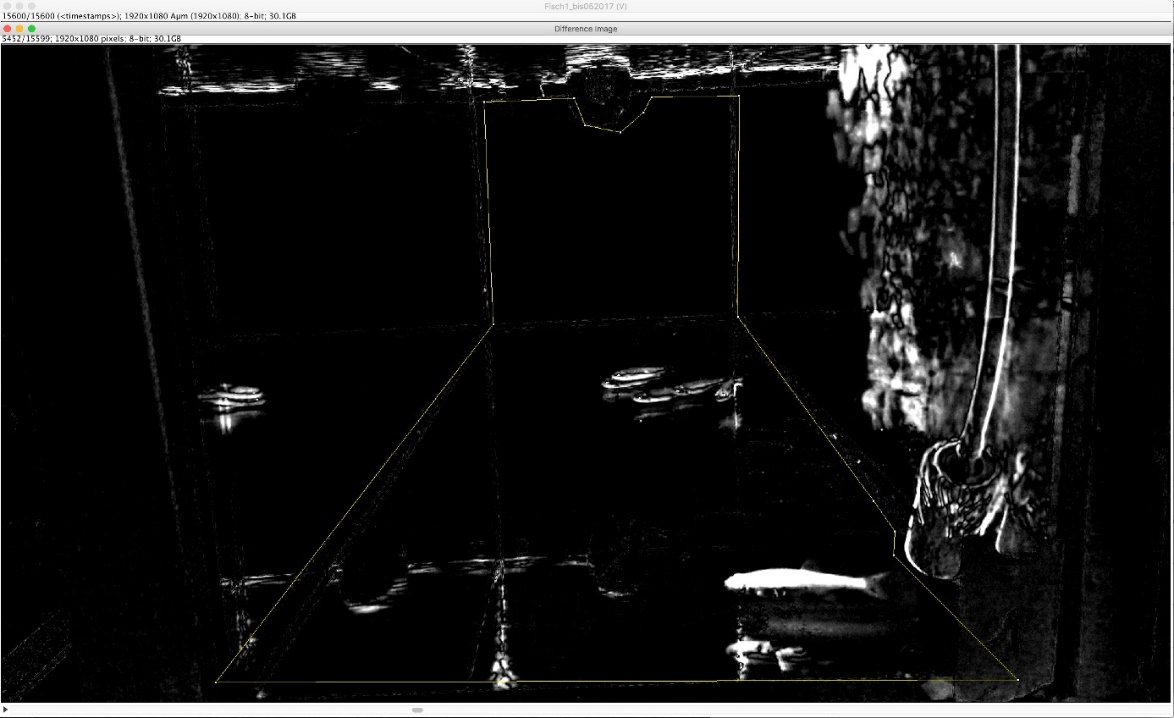

3. *Thymallus thymallus*

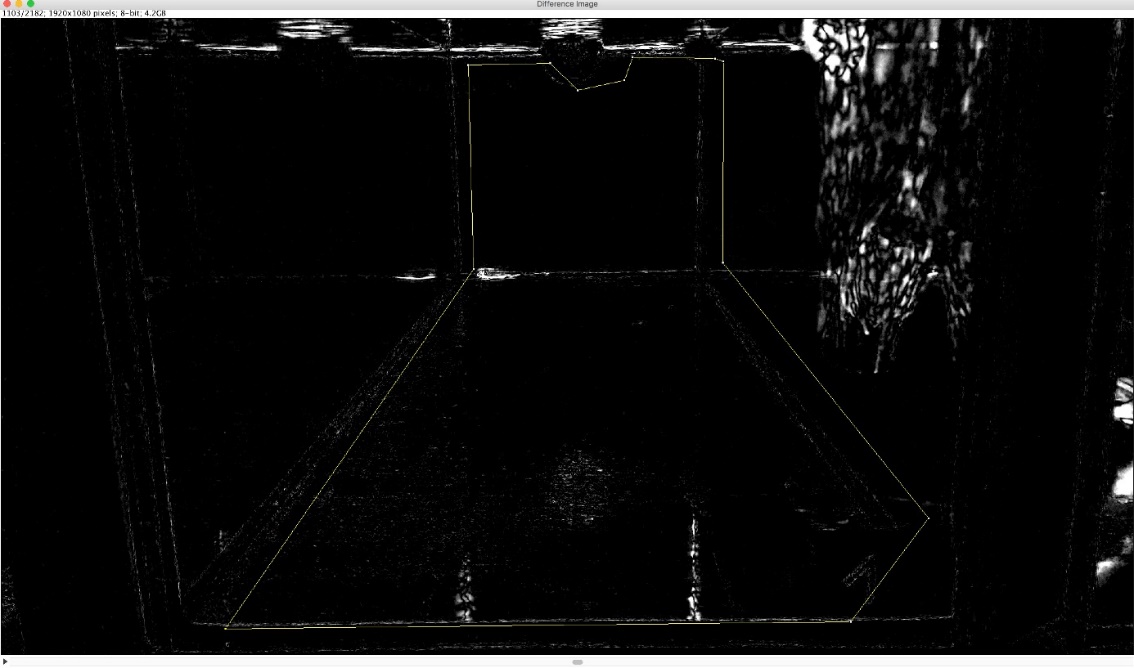

**Aquarium 2**

1. *Phoxinus phoxinus* single and group, *Squalius cephalus* single and group, *Oncorhynchus mykiss*

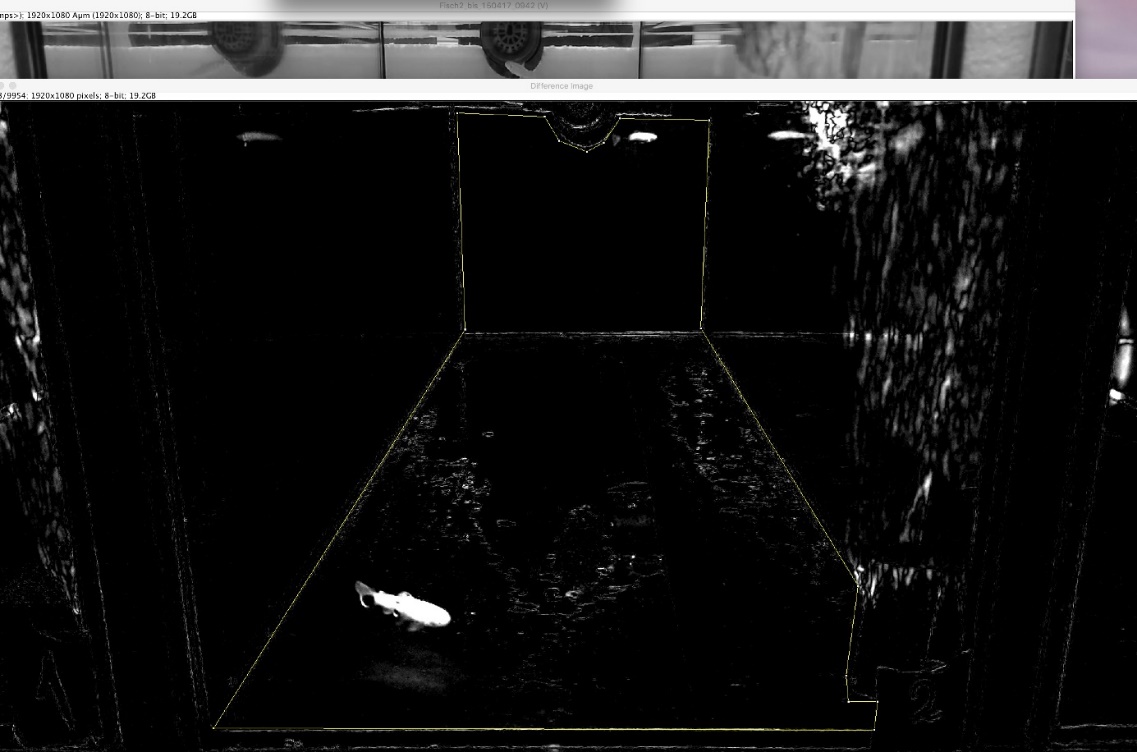

2. *Cottus gobio*, *Salmo trutta*, *Salvelinus fontinalis*

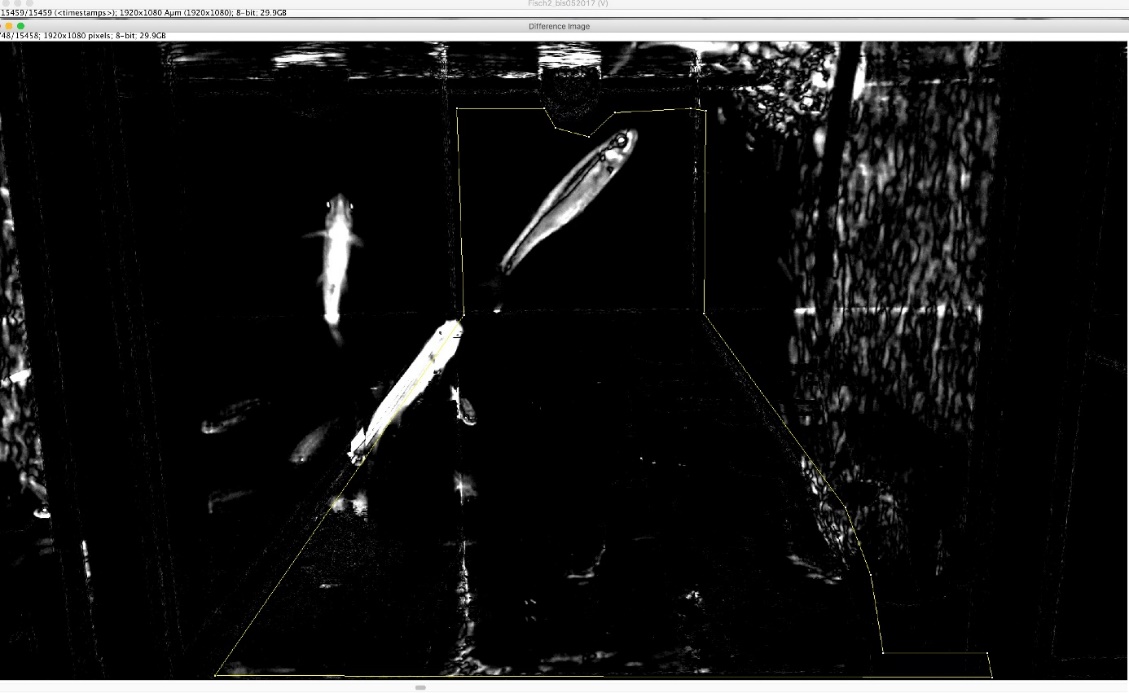

3. Beginning of *Thymallus thymallus*

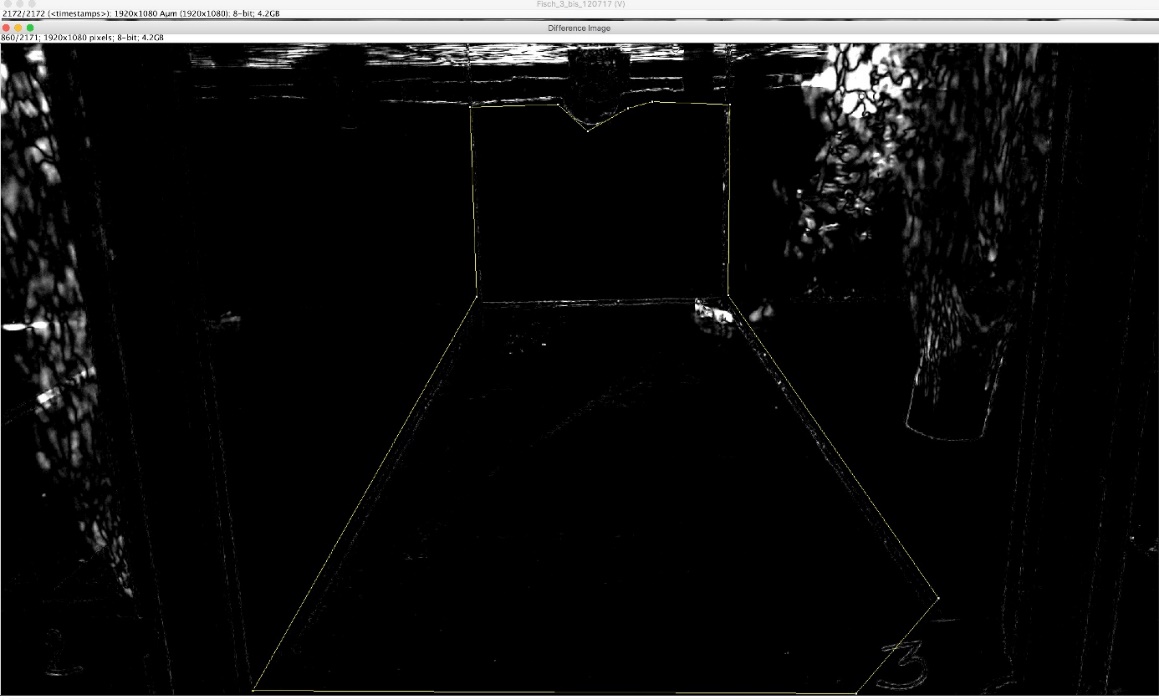

4. End of *Thymallus thymallus*

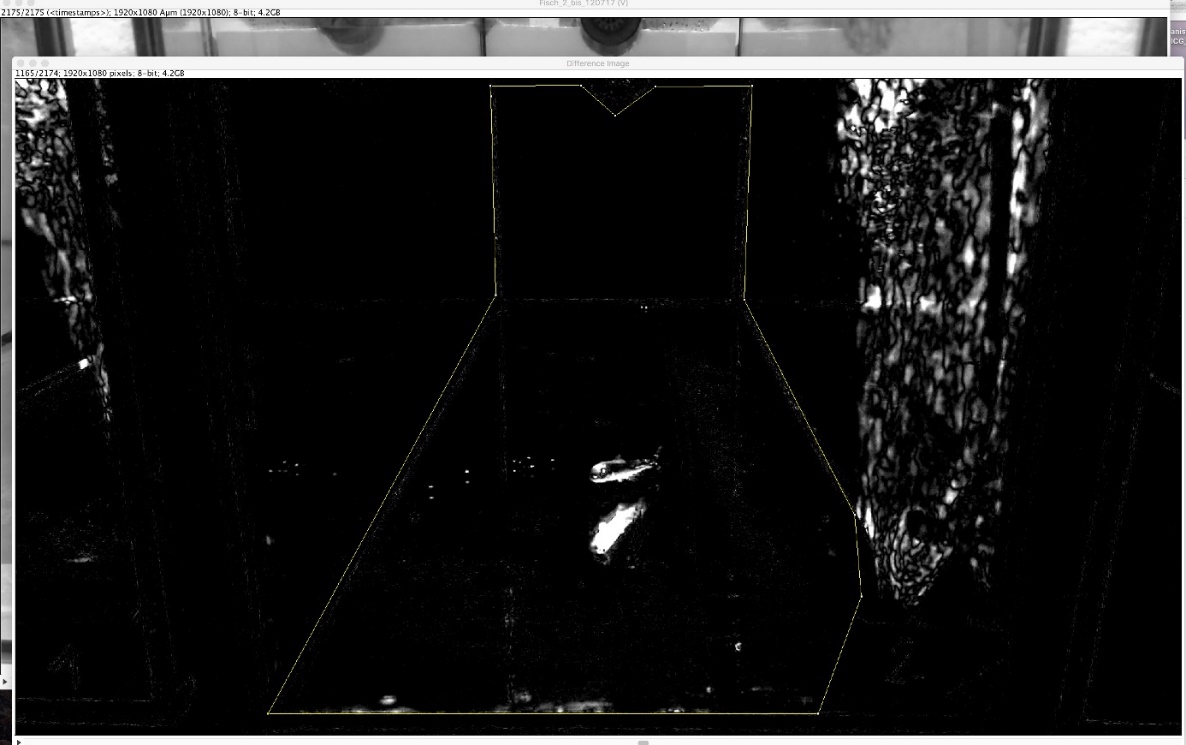

**Aquarium 3**

1. *Phoxinus phoxinus* single and group, end of *Squalius cephalus* single, *Squalius cephalus* group, *Oncorhynchus mykiss, Cottus gobio*, *Salmo trutta*, *Salvelinus fontinalis,*

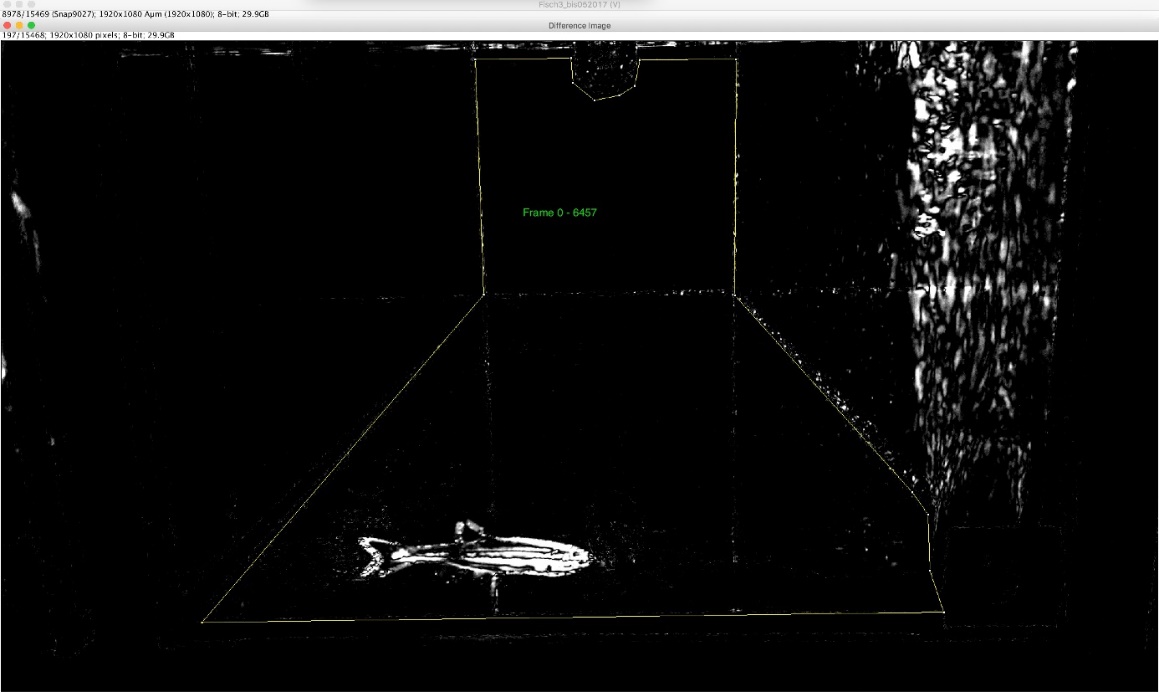

2. Beginning of *Squalius cephalus* single

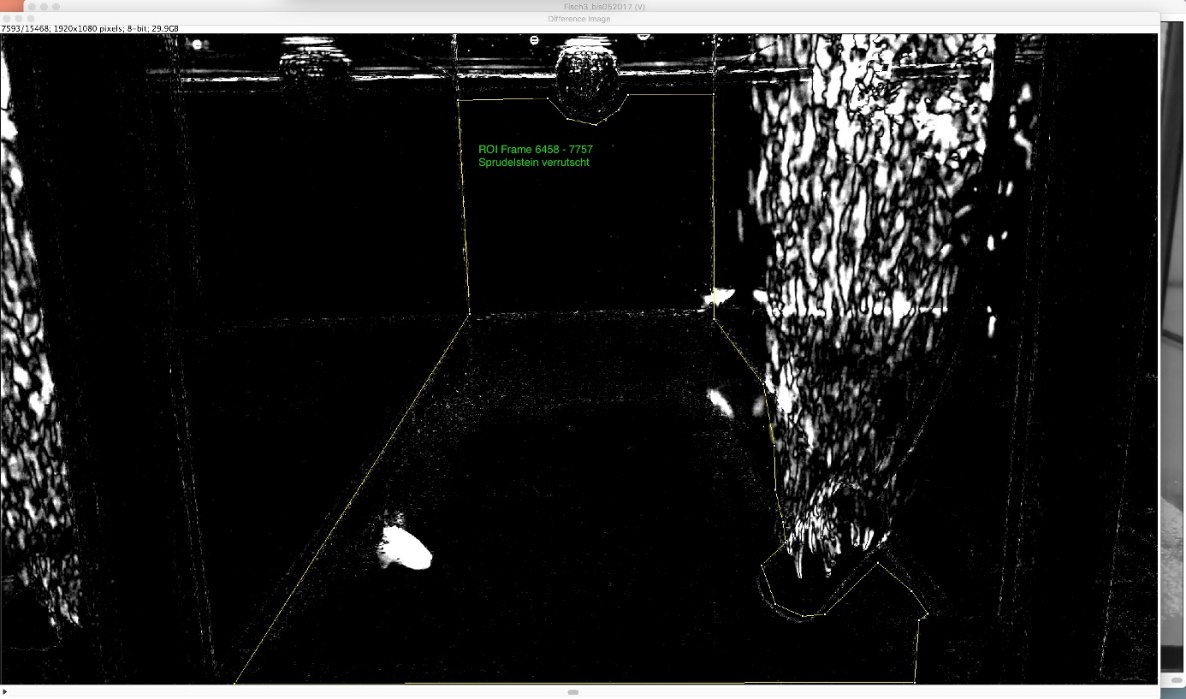

3. *Thymallus thymallus*

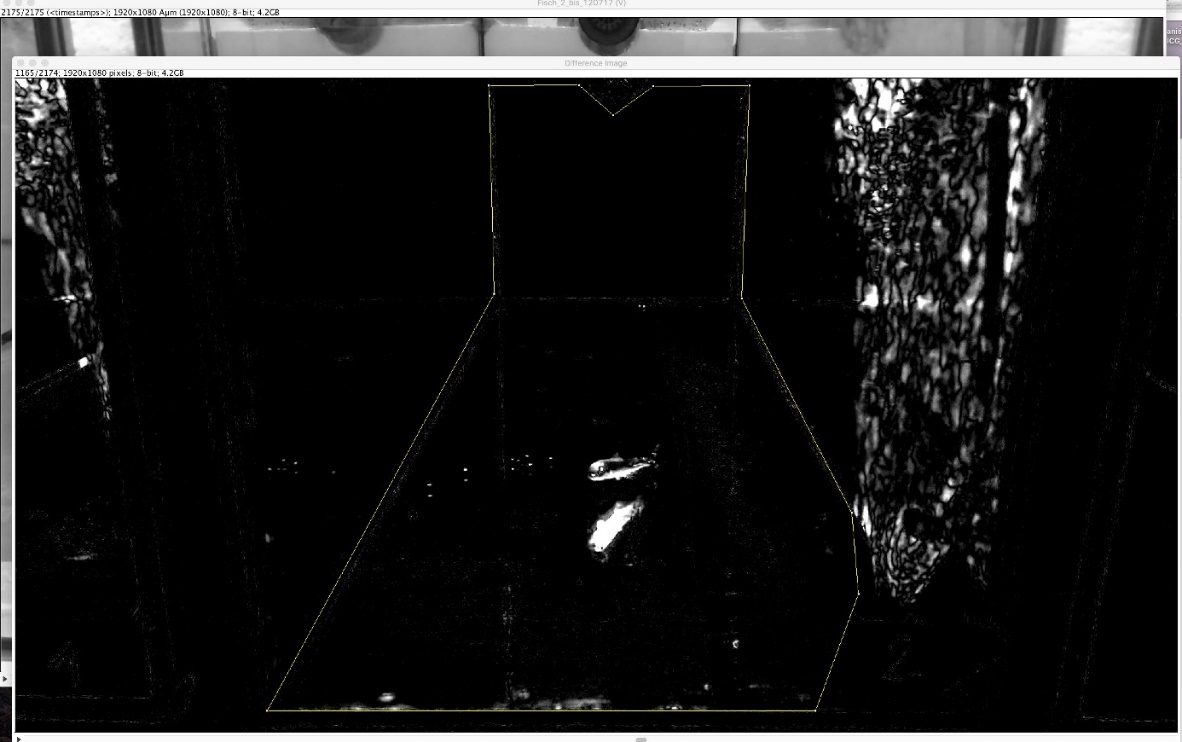

**Aquarium 4**

1. *Phoxinus phoxinus* single and group, *Squalius cephalus* single and group, *Oncorhynchus mykiss*

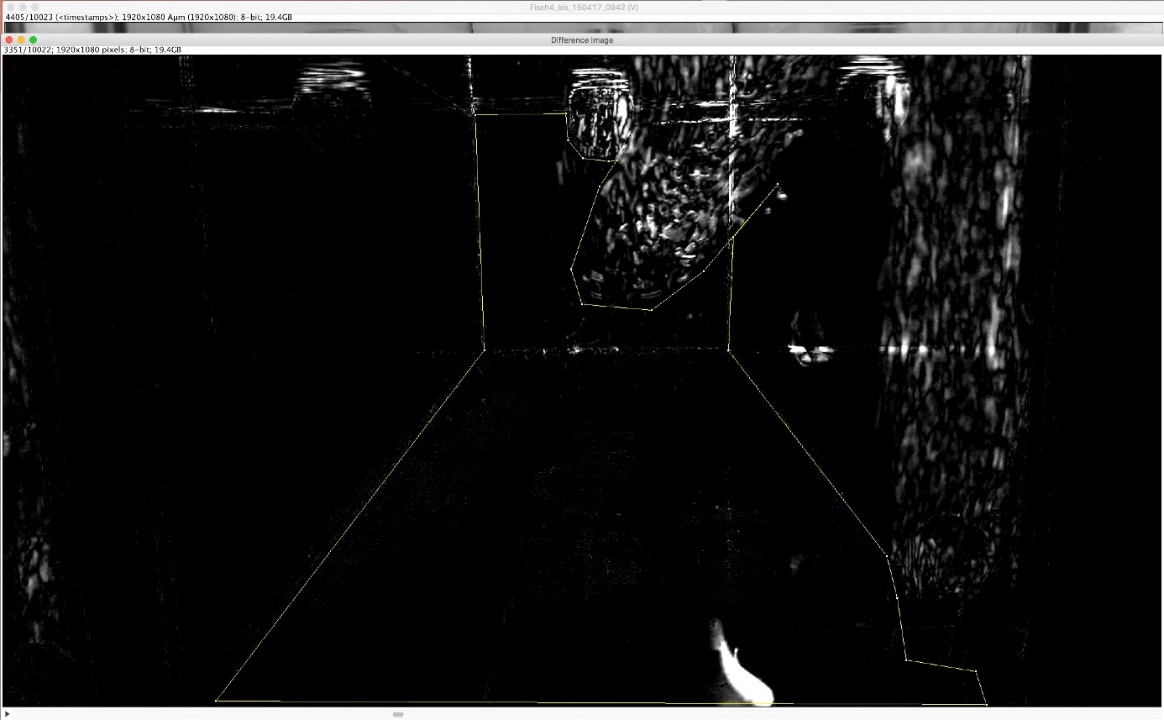

2. *Cottus gobio*, *Salmo trutta*, *Salvelinus fontinalis*

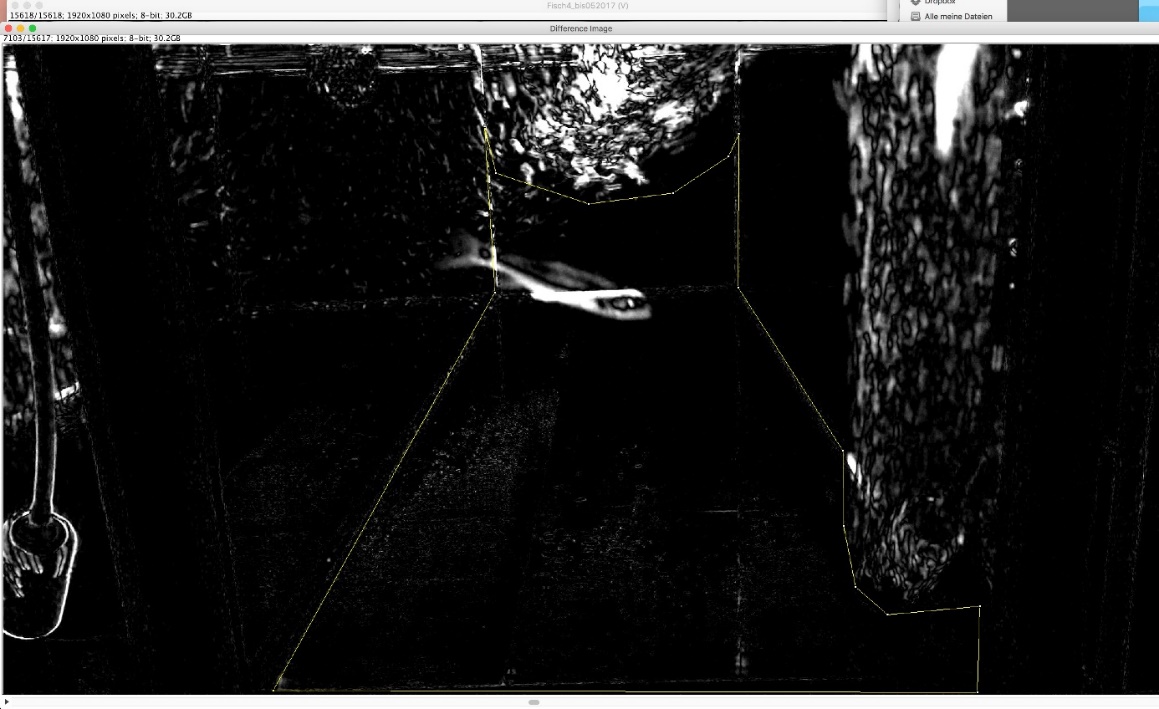

3. Beginning of *Thymallus thymallus*

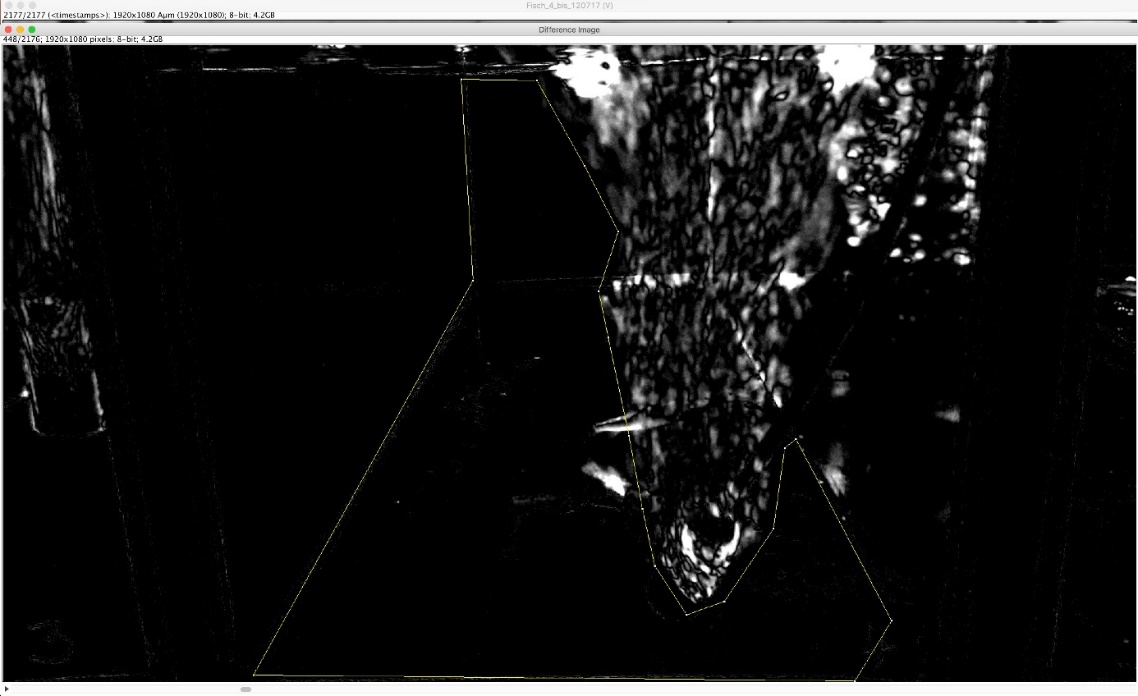

4. End of *Thymallus thymallus*

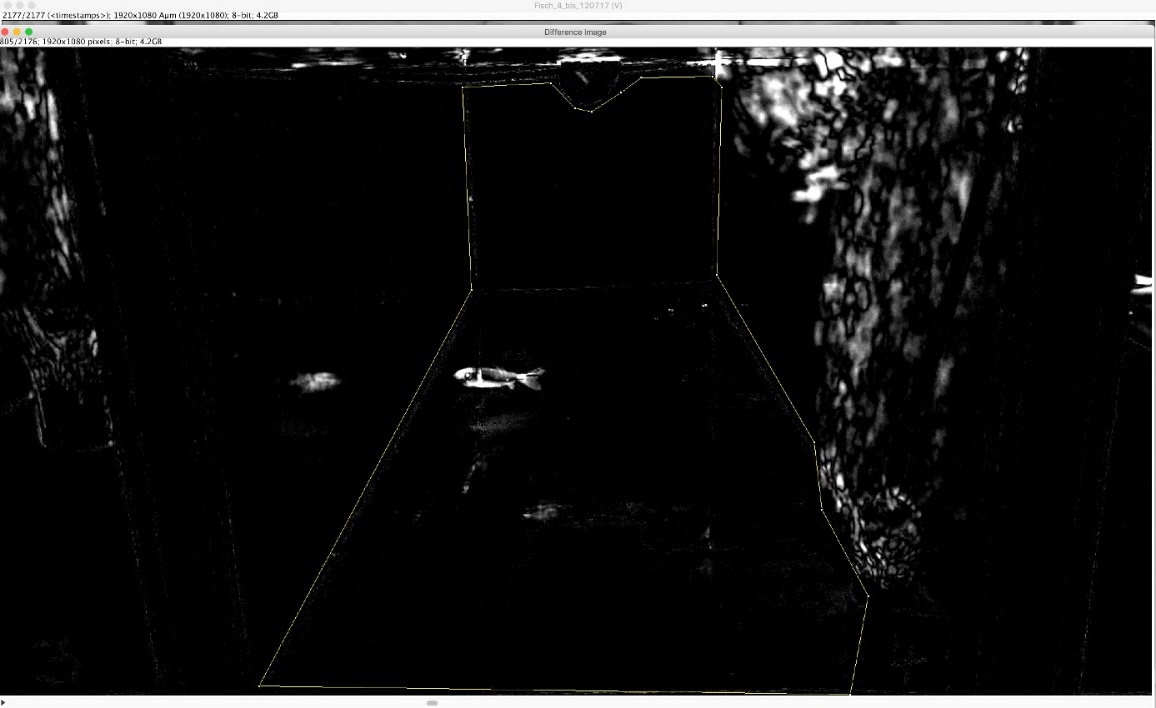

**Aquarium 5**

1. *Phoxinus phoxinus* single

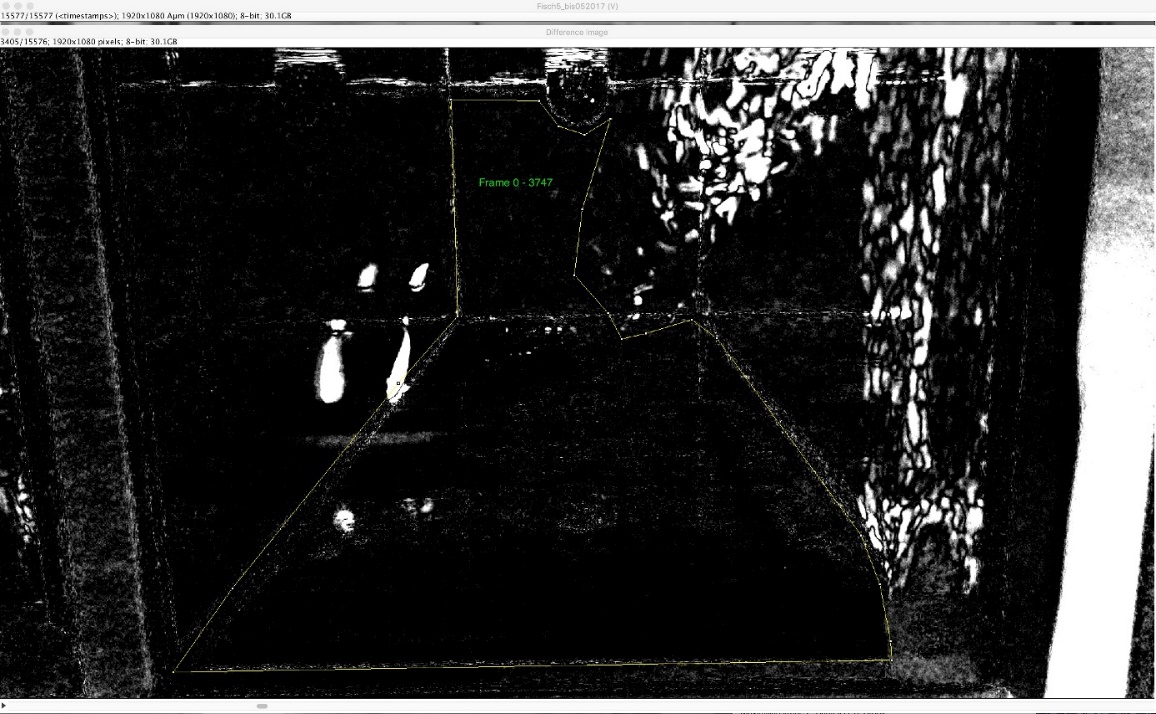

2. *Phoxinus phoxinus* group

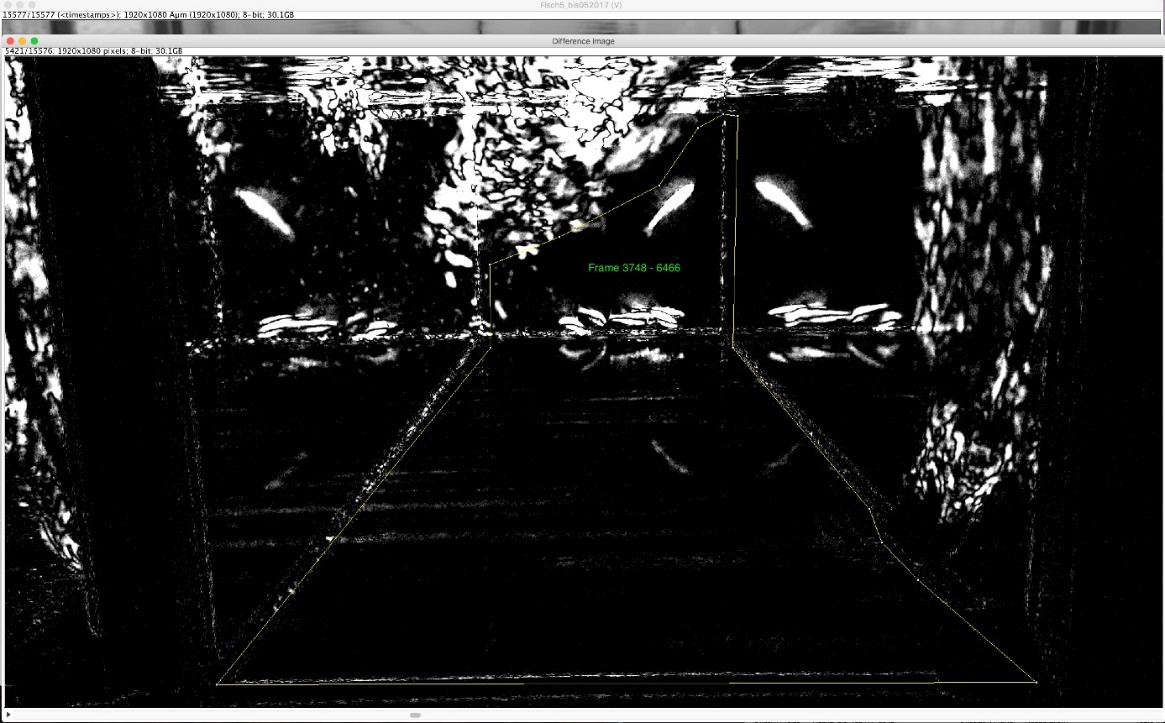

3. *Squalius cephalus* single, beginning of *Squalius cephalus* group

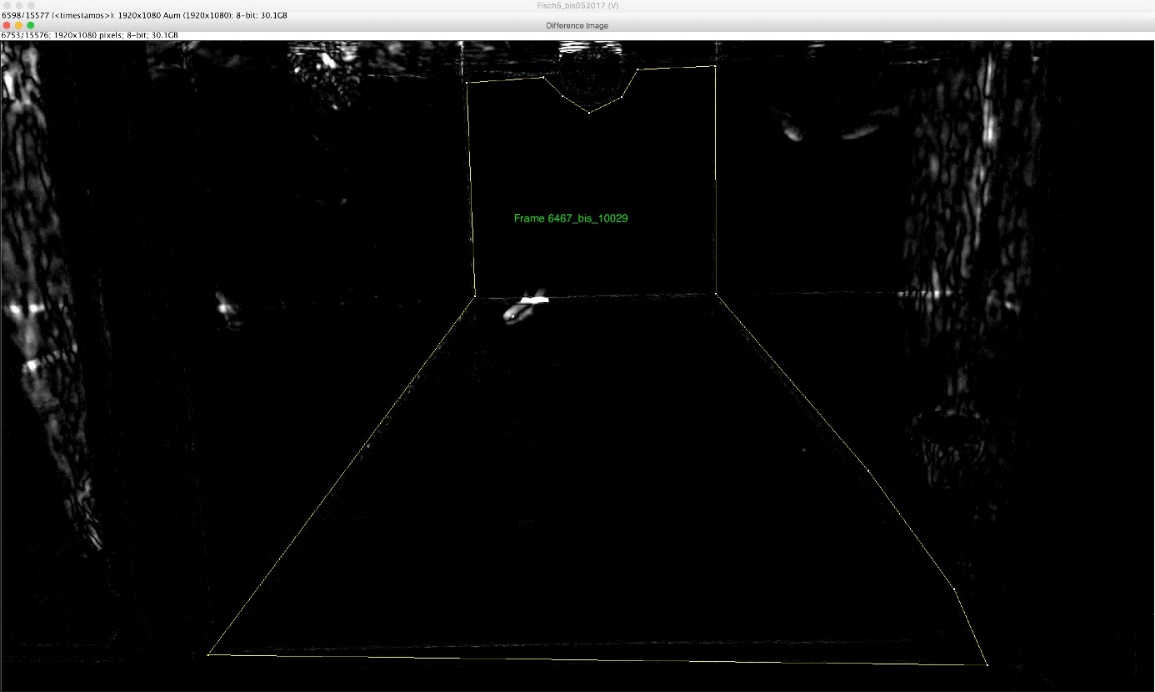

4. Middle of *Squalius cephalus* group

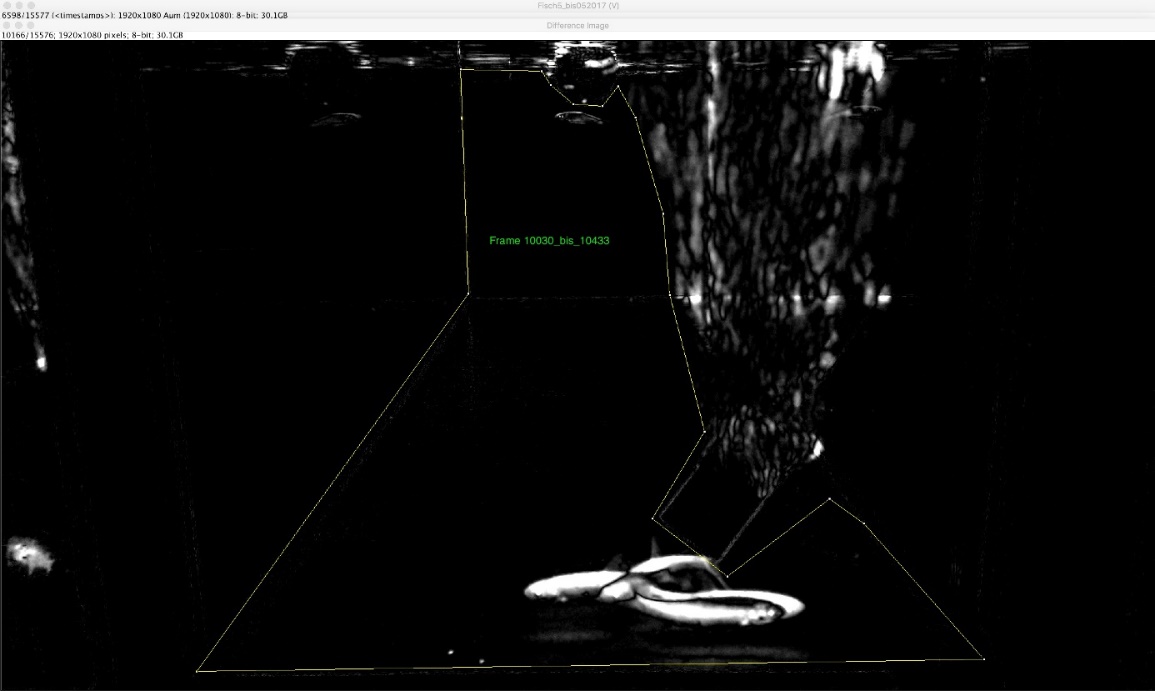

5. End of *Squalius cephalus* group, beginning of *Oncorhynchus mykiss*

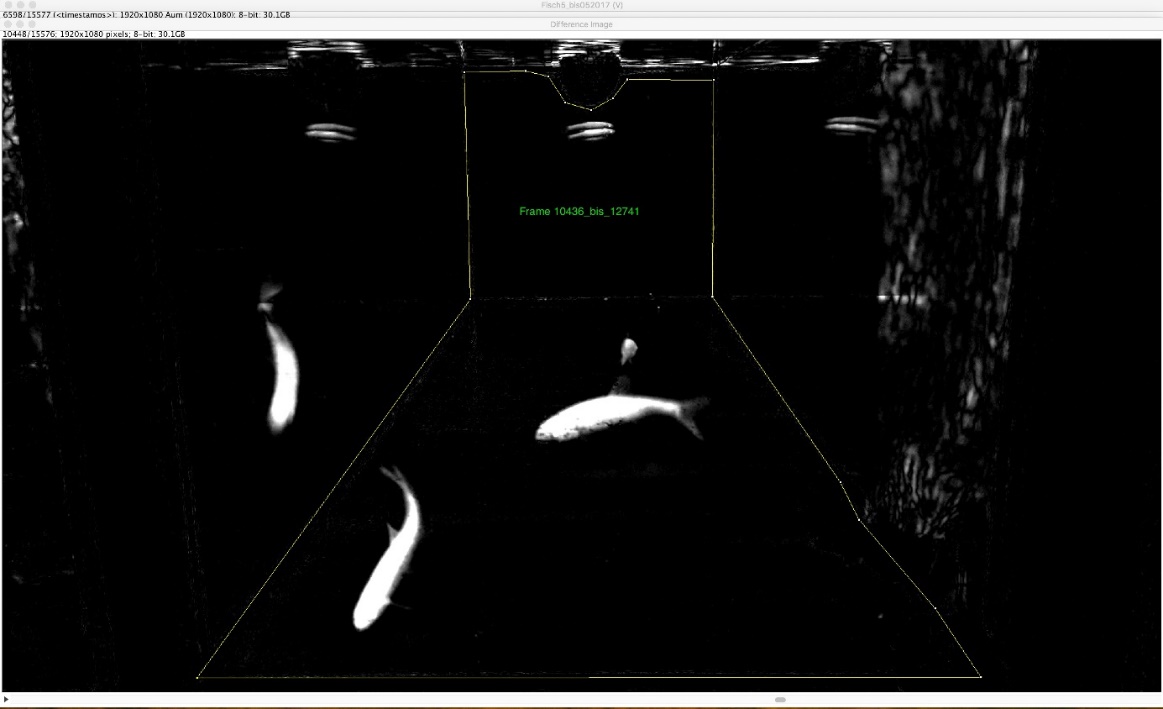

6. Middle of *Oncorhynchus mykiss*

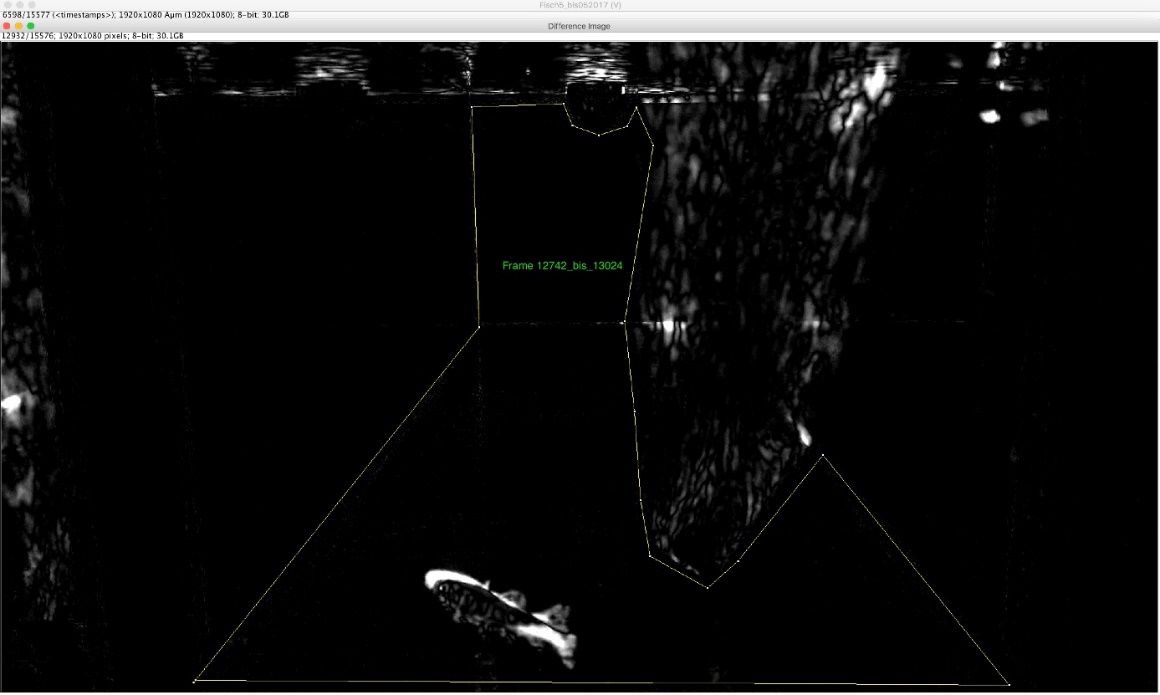

7. End of *Oncorhynchus mykiss*

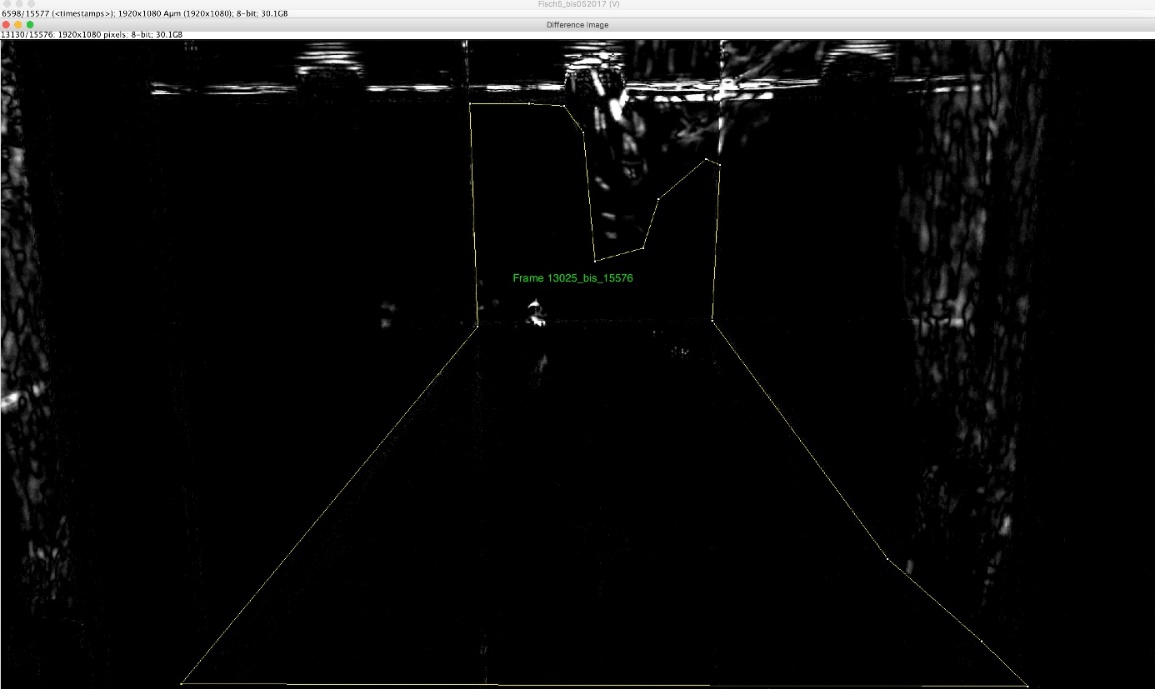

8. *Cottus cobio*

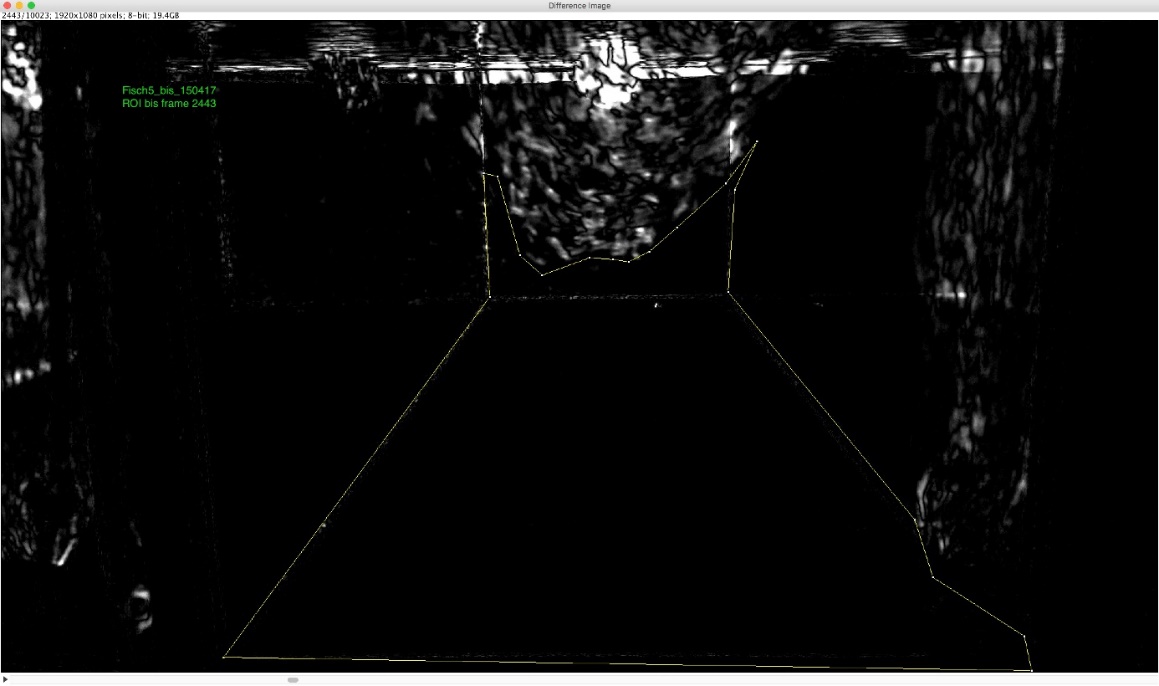

9. *Thymallus thymallus*

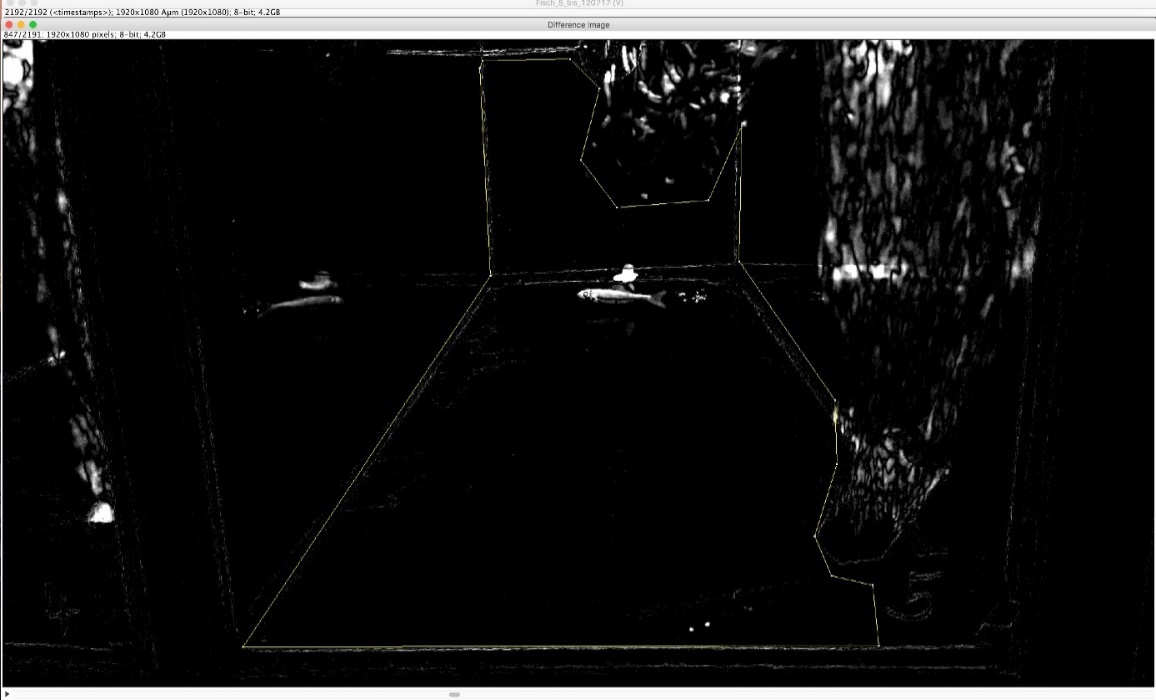

**Supplementary Material 3: Species-specific capillary electrophoresis PCR (celPCR)**

Prior to species-specific amplification via digital PCR (dPCR), all samples were subjected to celPCR to estimate their amount of fish DNA (Thalinger et al., 2019, 2020a, 2020b). DNA amplification was carried out with the Multiplex PCR Kit (Qiagen) in 10 µl PCRs including 3.2 µl DNA extract, one-time reaction mix, 5 µg BSA, 30 mM TMAC and 0.5 µM forward and reverse primer, respectively. The used primer pairs were previously validated for their specificity (Thalinger et al., 2016, 2020a). They reliably produce amplicons with a signal strength above 0.1 relative fluorescence units (RFU; see below) if at least 10 target DNA double strands are present at the start of PCR. Optimized thermo-cycling conditions were 15 min at 95°C, 35 cycles of 30 s at 94°C, 3 min at 64°C or 66°C (*P. phoxinus*), 1 min at 72°C and 10 min at 72°C once. For PCR product separation and analysis, the automated capillary electrophoresis system QIAxcel with the corresponding software QIAXCEL SCREENGEL version 1.4.0 (Method AM320; QIAGEN) was used. Samples resulting in target signals stronger than 0.08 RFU were deemed positive. The positive RFU values were noted and taken as a relative estimate of target DNA concentration in the samples (Thalinger et al., 2019, 2020a).

**Supplementary Material 4: The confounding variable fish mass**

During preliminary data analysis, fish mass was found to be a confounding variable, displaying a positive correlation with target eDNA copy numbers, mean activity and energy use (Fig. SM 4.1).

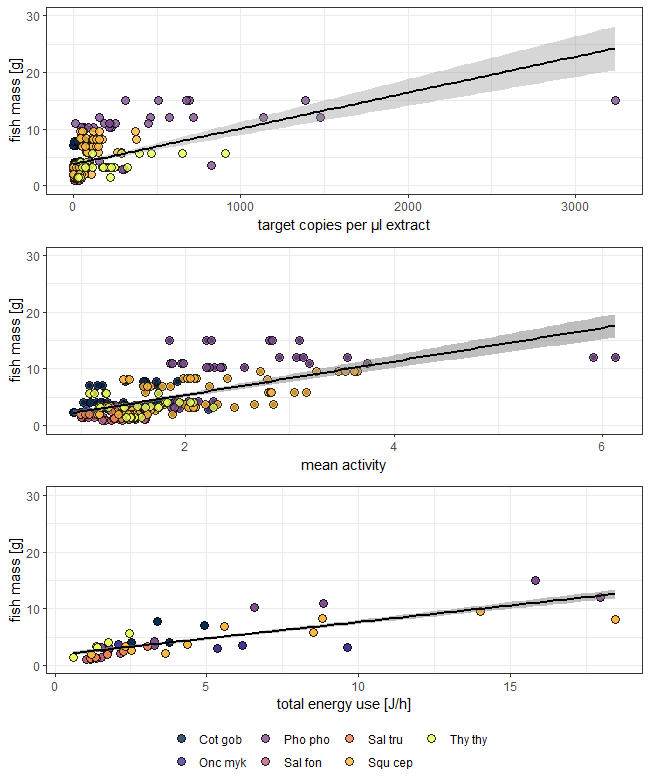

**Figure SM 4.1:** The positive relationship between fish mass and target eDNA copy number, mean activity and total energy use is displayed. Black lines depict the respective linear regression, grey areas the 95%CIs. Linear model equations from the top to bottom panel are y = 0.006*x + 3.79; R²-adj. = 0.29; y = -7.45*x + 8.71; R²-adj. = 0.46; y = 0.58*x + 31.81; R²-adj. = 0.65. Species are abbreviated as follows: “Cot gob“ = *Cottus gobio*, “Pho pho” = *Phoxinus phoxinus*, “Sal tru” = *Salmo trutta*, “Thy thy” = *Thymallus thymallus*, “Onc myk” = *Oncorhynchus mykiss*, “Sal fon” = *Salvelinus fontinalis*, and “Squ cep” = *Squalius cephalus.* Please note that measurements obtained from individual fish and fish groups were used for this analysis.

**References**

Thalinger, B., Kirschner, D., Pütz, Y., Moritz, C., Schwarzenberger, R., Wanzenböck, J., et al. (2020a). Lateral and longitudinal fish environmental DNA distribution in dynamic riverine habitats. *Environ. DNA* 00, 1–14. doi:10.1002/edn3.171.

Thalinger, B., Oehm, J., Mayr, H., Obwexer, A., Zeisler, C., and Traugott, M. (2016). Molecular prey identification in Central European piscivores. *Mol. Ecol. Resour.* 16, 123–137. doi:10.1111/1755-0998.12436.

Thalinger, B., Pütz, Y., and Traugott, M. (2020b). Endpoint PCR coupled with capillary electrophoresis (celPCR) provides sensitive and quantitative measures of environmental DNA in singleplex and multiplex reactions. *bioRxiv*. doi:10.1101/2020.10.24.353730.

Thalinger, B., Wolf, E., Traugott, M., and Wanzenböck, J. (2019). Monitoring spawning migrations of potamodromous fish species via eDNA. *Sci. Rep.* 9. doi:10.1038/s41598-019-51398-0.
